# Genomic and ecological factors shaping specialism and generalism across an entire subphylum

**DOI:** 10.1101/2023.06.19.545611

**Authors:** Dana A. Opulente, Abigail Leavitt LaBella, Marie-Claire Harrison, John F. Wolters, Chao Liu, Yonglin Li, Jacek Kominek, Jacob L. Steenwyk, Hayley R. Stoneman, Jenna VanDenAvond, Caroline R. Miller, Quinn K. Langdon, Margarida Silva, Carla Gonçalves, Emily J. Ubbelohde, Yuanning Li, Kelly V. Buh, Martin Jarzyna, Max A. B. Haase, Carlos A. Rosa, Neža Čadež, Diego Libkind, Jeremy H. DeVirgilio, Amanda Beth Hulfachor, Cletus P. Kurtzman, José Paulo Sampaio, Paula Gonçalves, Xiaofan Zhou, Xing-Xing Shen, Marizeth Groenewald, Antonis Rokas, Chris Todd Hittinger

## Abstract

Organisms exhibit extensive variation in ecological niche breadth, from very narrow (specialists) to very broad (generalists). Paradigms proposed to explain this variation either invoke trade-offs between performance efficiency and breadth or underlying intrinsic or extrinsic factors. We assembled genomic (1,154 yeast strains from 1,049 species), metabolic (quantitative measures of growth of 843 species in 24 conditions), and ecological (environmental ontology of 1,088 species) data from nearly all known species of the ancient fungal subphylum Saccharomycotina to examine niche breadth evolution. We found large interspecific differences in carbon breadth stem from intrinsic differences in genes encoding specific metabolic pathways but no evidence of trade-offs and a limited role of extrinsic ecological factors. These comprehensive data argue that intrinsic factors driving microbial niche breadth variation.

**One-Sentence Summary:** A nearly complete genomic catalog of the yeast subphylum illuminates the evolution of their diverse ecologies and metabolisms.

## Introduction

The ecological niche is a fundamental concept in ecology and evolutionary biology that explains the diversity and resource use of organisms through space and time. Interest in how niche breadth evolves dates to Darwin, and studies of niche breadth evolution encompass diverse topics that include the evolution of species diversity, trait coevolution, and the assembly of communities. Species with broad niche breadths are defined as generalists, while those with narrow ones are specialists. There are many biotic and abiotic dimensions of the niche that can vary among organisms (*1–3*). The presence of extensive variation in niche breadth begs the question: Why do both generalists and specialists exist? In other words, what underlying genomic and ecological factors contribute to niche breadth variation?

Two broad paradigms have been offered as answers. The first is that niche generalism and specialism are governed by trade-offs between performance efficiency and breadth (*4–9*). For example, replicate populations of *Escherichia coli* grown in medium containing only glucose repeatedly evolved increased ability to use glucose but showed a reduction in their ability to catabolize other carbon sources (*10*). In contrast, strains of *Saccharomyces cerevisiae* grown in glucose limitation conditions gained fitness advantages across multiple sugar-limitation environments (*3*), while positive correlations between thermal tolerance ranges and immune responses were detected across several major clades of endothermic and ectothermic vertebrates (*11*). These numerous divergent reports question the generality and even the existence of trade-offs (*3*, *5*, *10–21*).

The second paradigm suggests that diverse intrinsic and extrinsic factors drive variation in niche breadth. Intrinsic factors include the evolution of promiscuous enzymes responsible for the utilization of multiple resources (*22–26*), as well as overlapping biochemical, developmental, or genetic pathways (*27*, *28*). For example, *MAL* and *IMA* genes are promiscuous enzymes associated with the utilization of multiple carbon sources in yeasts, that is they can increase niche breadth by enabling broader consumption (*22*, *23*). Conversely, gene loss due to drift or relaxed selection could lead to narrower niche breadth (*29*). Niche breadth itself, therefore, is not the primary trait under selection; instead, niche breadth is the sum of evolutionary forces acting on individual intrinsic traits. Extrinsic factors, such as environmental heterogeneity, could also exert selective pressure on traits, resulting in generalism and specialism (*30*). Environments with high levels of spatial or temporal heterogeneity can favor generalists, while homogeneous environments favor the evolution of specialists (*31–37*).

The subphylum Saccharomycotina of kingdom Fungi, which includes the model baker’s yeast *Saccharomyces cerevisiae*, the opportunistic pathogen *Candida albicans*, and the oleochemical cell factory *Yarrowia lipolytica*, exhibits extensive ecological, genomic, and metabolic diversity; thus, it is a superb system for testing the two major paradigms for the evolution of metabolic niche breadth (Fig. 1). Species in the subphylum, commonly referred to as yeasts, exhibit extensive genomic diversity; levels of gene sequence divergence across yeasts are comparable to levels observed across plants and animals, and the subphylum also harbors considerable variation in gene content, including metabolic genes (*22*). In addition, extensive experimental work in model yeasts, such as *S. cerevisiae* (*38*) and *C. albicans* (*39*), provides validated functional genetic information. Yeast growth profiles have been characterized across many carbon and nitrogen sources and environmental conditions (e.g., temperature), and they are highly variable across species (*22*, *23*, *40*). This phenotypic diversity is a major driver of yeast ecological diversity. Yeasts are found in nearly every biome of the world on a wide array of substrates, and the isolation environments of these yeasts are associated with specific phenotypic traits. For example, both glucose and sucrose fermentation are positively associated with fruits, fermented substrates, and juices (*23*). This positive association is driven by multiple genera of yeasts that have been linked to wine production and food spoilage (*23*, *41*, *42*). This treasure trove of genomic, metabolic, and environmental diversity across an entire subphylum makes Saccharomycotina an attractive and highly tractable system for studying niche breadth evolution.

**Figure 1:**
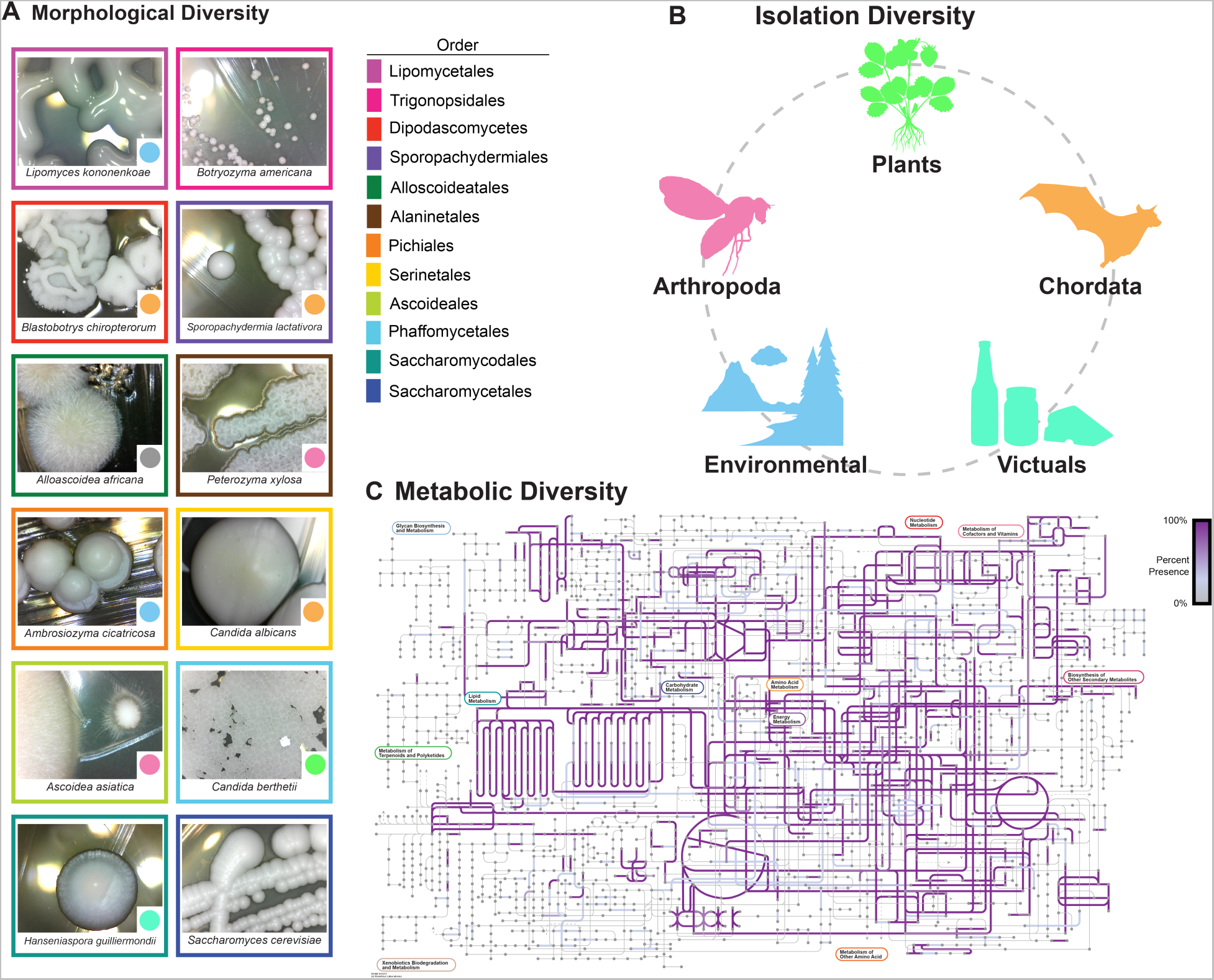
Yeasts are morphologically, ecologically, and metabolically diverse. **A.** Yeast colonies are morphologically diverse; they can vary in shape, color, size, dullness, etc. Images of yeasts from different orders. The color of the box surrounding the image indicates the species’ order. The color of the circle in the bottom right-hand corner of the image represents the isolation environment for the strain of the species sequenced and phenotyped during this study. **B.** Yeasts have been isolated from every biome and continent. Strains studied were found on plants, animals, in soil, and many other environments. **C.** Yeasts are metabolically diverse. The image represents the KOs present across Saccharomycotina metabolic networks. Any pathway that is highlighted in purple is present across a subset of yeasts; the saturation of the purple represents the proportion of yeasts with the pathway.

To study the evolution of generalists and specialists, we quantified variation in genome sequence, isolation environment, and carbon and nitrogen metabolism for 1,154 yeast strains, which represent nearly all known species in the subphylum Saccharomycotina. We generated draft genomes for 1,086 strains, which includes 1,016 described species and 62 candidates for novel species, and measured the quantitative growth of 853 strains on 24 carbon and nitrogen sources over the course of the Y1000+Project (http://y1000plus.org) (*43*). Yeasts were assigned to a niche breadth classification—specialist, standard, or generalist—for both carbon and nitrogen metabolism. This novel and rich dataset enabled us to test multiple hypotheses of niche breadth evolution, including trade-offs between breadth and growth rate, coevolution between traits, and whether intrinsic (i.e., genomic) and extrinsic (i.e., isolation environment) factors drive the evolution of generalists and specialists. Our analyses of niche breadth evolution across an entire subphylum find no evidence of trade-offs, rejecting the “jack-of-all-trades, master of none” paradigm, and do not support a major role for extrinsic factors. Rather, our machine learning analyses pinpoint specific genetic differences between generalists and specialists, including novel associations between carbon generalism and specific metabolic pathways, suggesting that intrinsic factors are the primary drivers of yeast niche breadth variation.

## Results and Discussion

### A genomic, evolutionary, and metabolic portrait of a eukaryotic subphylum

We sequenced and assembled 953 new genomes in this study and combined them with 140 genomes previously sequenced by the Y1000+ Project and 61 publicly available genomes (table S1A). Our dataset contained 1,154 genomes from 1,049 species, including 1,034 taxonomic type (i.e., ex-type) strains of currently recognized species. Multiple strains were sequenced for 42 species, including 8 recognized varieties. Sixty-two candidates for new species were also sequenced. The genomic dataset spans 96 yeast genera, which is 95% of the currently described genera. The genera not included are those for which no living culture is available (i.e., *Endomyces, Coccidiascus, Helicogonium, Phialoascus*, and *Macrorhabdus*), as well as genera described after our last round of genome sequencing in February 2021 (*Limtongella, Babjevia, Hemisphericaspora, Metahyphopichia*, and *Savitreea*). Our genome sequencing added between 1 and 336 species to each order, most notably expanding the order Serinales (previously major clade CUG-Ser1), which contains the human pathogens *C. albicans* and *Candida auris*, from 94 genomes to 430. All genome assemblies totaled ~15 billion base pairs. The assemblies had a mean N50 of 387.5 Kb, which is comparable to our previous smaller scale dataset of 332 genomes (417.2 kb) (*22*). Of the 1,154 yeast genomes, 1,000 (~87%) have ≥ 90% of the 2,137 predefined single-copy orthologs defined by OrthoDB v10 (*44*, *45*) (table S1A & table S1B).

To infer the genome-scale phylogeny of the Saccharomycotina, we used 1,403 orthologous groups (OGs) from 1,154 Saccharomycotina genomes and 21 outgroups. Nearly all internodes in both concatenation-based (1,136 / 1,153; 99%) and coalescent-based (1,123 / 1,153; 97%) phylogenies received strong (≥ 95%) support (Fig. 2 and fig. S1 and fig. S2). The two phylogenies were highly congruent, with only 60 / 1,153 (5%) conflicting internodes (fig. S2). Moreover, relationships among the 12 recently circumscribed taxonomic orders (*46*) (previously major clades) were congruent with previous studies (*22*, *47*, *48*), including the placement of the Ascoideales (previously CUG-Ser2) and Alaninales (previously CUG-Ala).

**Figure 2:**
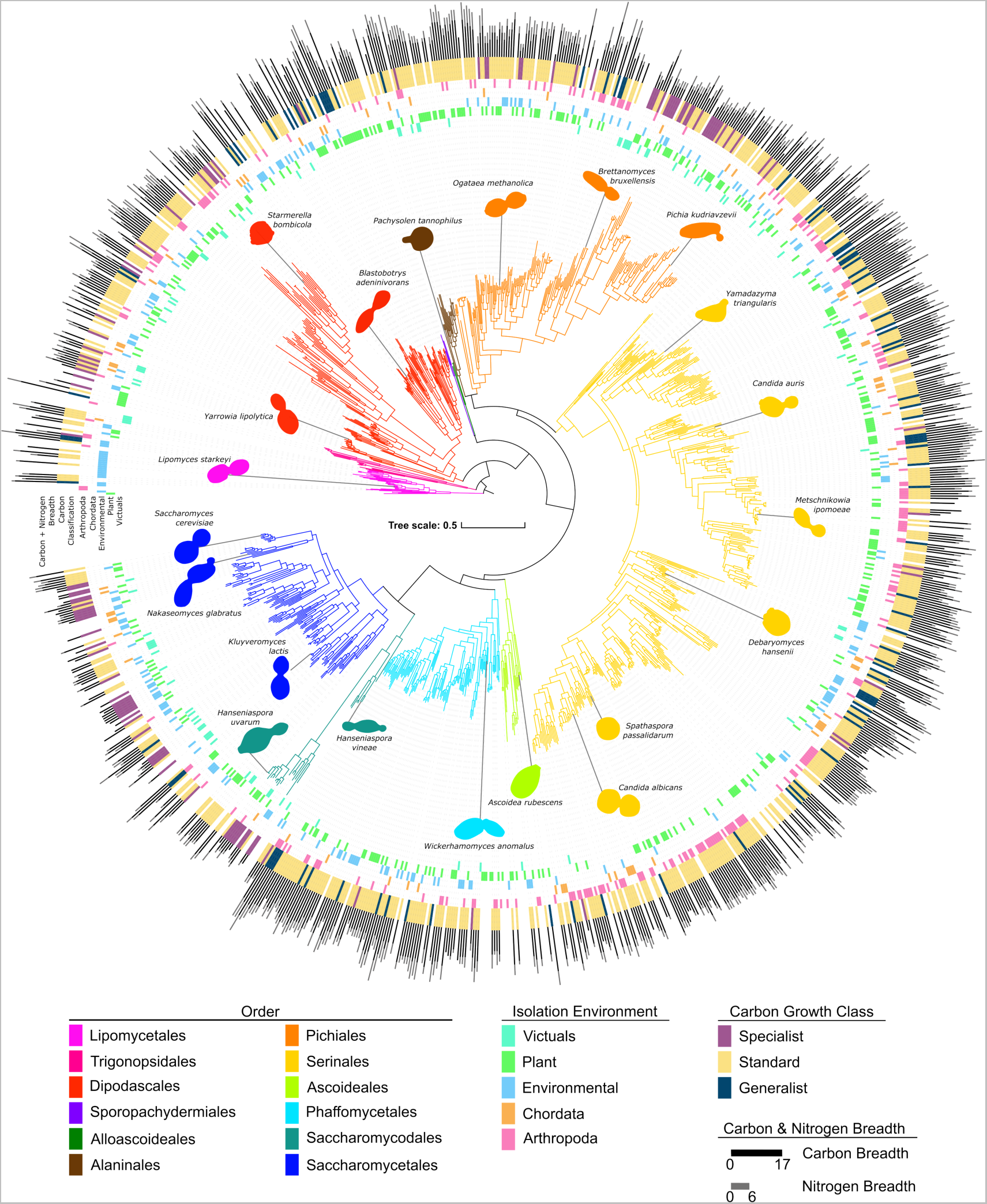
Yeast traits are widely distributed across the phylogeny. The phylogeny of 1,154 yeasts and fungal outgroups built from 2,408 orthologous groups of genes. Branches are colored according to their taxonomic assignment to an order of Saccharomycotina (*46*). The innermost rings are colored by the top-level type of isolation environment in which each specific strain was isolated. The purple, yellow, and blue ring identifies the carbon growth classification for each strain. This classification is based on the carbon breadth, which, along with nitrogen breadth, is represented by the bar graph on the exterior of the tree. All of the traits illustrated (isolation environment, carbon growth class, nitrogen breadth, and carbon breadth) are widely distributed across the tree; no order has one trait exclusively.

Analyses of the inferred codon tables and tRNA genes (Figshare repository) were consistent with previous codon reassignments. Notably, the Ascoideales had a diversity of tRNAs with CAG anticodons predicted to decode CUG codons. The Ascoideales CAG anticodon was associated with multiple isoreceptors across species: leucine (n=1), leucine and serine (n=9), serine (n=6), and valine and serine (n=5). Despite a predicted CAG-tRNA with a valine isotype, a computational screen of gene sequences did not predict a novel reassignment. The diversity of tRNA species in the Ascoideales is consistent with previous findings that its member species may stochastically decode CUG as both leucine and serine (*49*).

To examine the evolution of metabolic niche breadth across Saccharomycotina, we quantified the growth rates of 853 yeast strains on 17 carbon sources, 6 nitrogen sources, and a no carbon control (table S2). We found that yeasts displayed variation in growth rates across carbon (fig. S3A) and nitrogen sources (fig. S3B); on average, yeasts could metabolize eight carbon (fig. S4A) and two nitrogen sources (fig. S4B). Comparison of growth rates on different carbon sources revealed that 65.22% of yeasts (n = 557) grew fastest on glucose, with the remaining 34.78% (n = 297) growing faster on a non-glucose carbon source (fig. S5). Mannose, an epimer of glucose not typically tested in yeast growth experiments, was the carbon source that yeasts grew fastest on after glucose (n = 112). We also found that 77 yeasts grew faster on fructose than glucose, including cases where their maximum growth rate was on a third carbon source. Several of these yeasts (n = 7) were in the *Wickerhamiella/Starmerella* clade, which contains many known fructophilic yeasts (*50*). The fructophilic results were independently verified in a second lab on a subset of yeasts (table S3). Together, these results challenge the widely held assumption that glucose is the preferred carbon source of all yeasts.

### No trade-offs between carbon breadth and carbon growth rates

We classified yeasts into three categories for both carbon and nitrogen utilization niche breadths: specialist, standard, and generalist (table S2). We found that, for both carbon and nitrogen metabolism, most yeasts were standard (75.9%: 648/853 and 78.4%: 669/853, respectively) (table S2). From the remaining 24.0% (n = 205/853) of carbon generalists and specialists, 54.3% (n = 110/205) were specialists, and 46.3% (n = 95/205) were generalists (Fig. 3A). The median numbers of carbon sources used by specialists and generalists were four and twelve, respectively. Carbon generalists and specialists were widely distributed across the subphylum (Fig. 2), and orders with more than 15 phenotyped strains (n = 8) all featured both generalists and specialists. However, the relative proportion of generalists and specialists within orders varied greatly. For example, the Saccharomycetales (n=82) had 3 generalists and 33 specialists, while the Serinales (n= 347) had 53 generalists and 9 specialists. This result suggests that yeast orders exhibit distinct eco-evolutionary trajectories.

**Figure 3:**
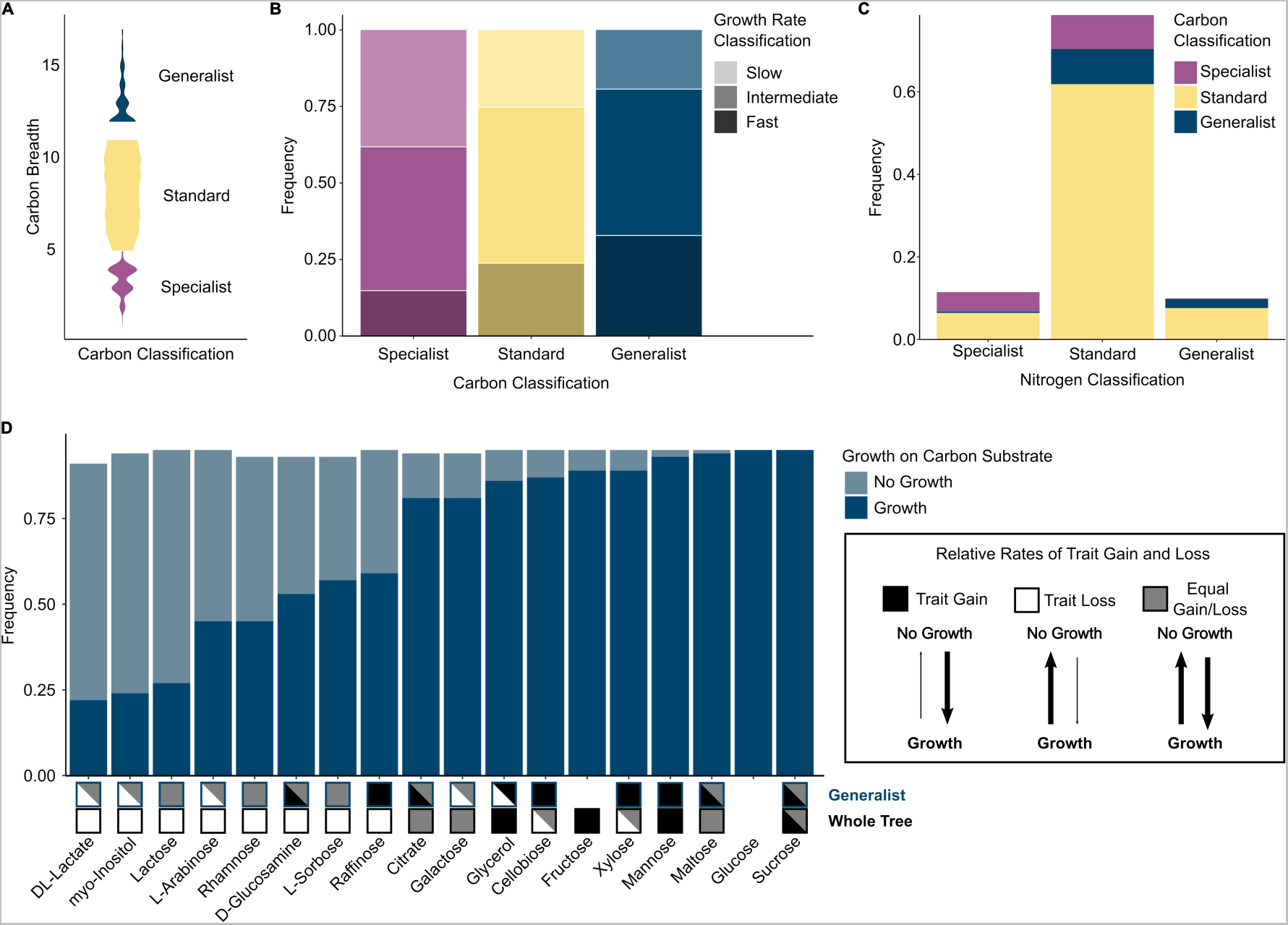
Carbon specialists and generalists differ in nitrogen breadth, growth rate, and evolutionary history. **A.** Violin plot of carbon breadth for each carbon metabolic strategy across 853 yeast strains. Metabolic classifications were determined by permuting the binary carbon growth matrix (n = 1000 permutations). To determine the metabolic strategy of a species, we calculated the observed and expected (permuted) breadth for each species and calculated the binomial confidence intervals to determine significant differences in breadth. Generalists have a significantly larger carbon breadth than expected by chance, and specialists have a significantly smaller carbon breadth. If a species was not classified as either a generalist or a specialist, it was classified as standard. **B.** The growth rates for each species on each of the 17 carbon sources were categorized as slow (bottom 25%), intermediate (median 50%), or fast (top 25%). Carbon generalists had the highest proportion of fast growth rates (33%), while specialists had the smallest proportion (15%.) The inverse was also true, with carbon generalist having the smallest proportion of slow growth rates (19%) and carbon specialists having the highest proportion of slow growth rates (38%). **C.** Stacked bar graph of carbon metabolic strategies within each nitrogen metabolic strategy. Across all three nitrogen metabolic strategies, most yeasts had a standard carbon metabolic strategy. **D.** Carbon generalists shared many of the same growth traits – 10 out of 17 growth traits were found in more than 75% of generalists. Many of the carbon sources had different evolutionary trends in a generalist background as compared to across the whole tree. Three different evolutionary models are shown: trait gain (black), trait loss (white), and equal rates of trait gain and loss (gray). No box indicates that the trait was not co-evolving with background or across the tree. More than one evolutionary model is shown in cases where the reverse jump model spent 75% or less of the time on a single model. For example, the model testing correlated evolution between growth on D-glucosamine and generalist carbon classification reported a model string with a greater rate of gain in 55% of the run and a model string with equal rates of gain and loss in 29% of the run; therefore, we reported both the trait gain and equal gain/loss model in the generalist analysis.

We tested for a trade-off between growth rate and carbon breadth by classifying growth into three categories: slow (growth rate in the lower quartile), intermediate, and fast (growth rate in the upper quartile.) Within each carbon source, we found specialists were more often slow growers (38%: 146/381 growth rates) than fast growers (15%: 54/381) (Fig. 3B). In contrast, generalists were more often fast growers within each carbon source (33%: 403/1222 growth rates), rather than slow growers (20%: 238/1,222 growth rates) (Figure 3B). Moreover, none of the tested carbon sources had more specialist fast growers than generalist fast growers (table S4). We also examined linear phylogenetically corrected correlations between growth rates and carbon breadth. We found that growth rates on four carbon sources (glucose (FDR p = 0.0028), mannose (FDR p = 0.0056), myo-inositol (FDR p = 0.0083), and fructose (FDR p = 0.0111)) were positively correlated with carbon breadth when accounting for phylogeny (table S5). No negative correlations were identified, which would have indicated that specialists were faster growers. Our finding that generalists grew faster on more substrates than specialists suggests that, at least under laboratory conditions, there is no trade-off between carbon breadth and carbon growth rate.

We next tested whether there is a trade-off between carbon and nitrogen breadth. Relative to all other combinations (e.g., nitrogen generalist and carbon specialist), we found significantly fewer carbon generalists that were nitrogen specialists (n = 1) and carbon specialists that were nitrogen generalists (n = 2) than statistically expected (χ^2^= 69.25, p-value = 3.26×10^−14^) (Fig. 3C). Moreover, evolutionary analysis using BayesTraits found that carbon and nitrogen generalism coevolve (Bayes factor >2; full results in Figshare repository). The models of dependent evolution found nitrogen generalism in a carbon specialist/standard background evolved at a rate of zero or close to zero, suggesting that nitrogen generalism arises almost exclusively in a background of carbon generalism (table S6). In other words, carbon generalism mainly arises before and may be required for nitrogen generalism. Additionally, phylogenetic regression analysis showed a strong positive correlation between carbon breadth and nitrogen breadth (reported p-value of 0.000, slope of correlation = 0.92; table S6). Rather than a trade-off, these results suggest that there is an evolutionarily conserved functional connection between carbon and nitrogen metabolism in yeasts. Consistent with our finding, it is well known that certain amino acids can serve as both a carbon and nitrogen source and, as such, are dually regulated by both carbon and nitrogen signaling systems (*51*, *52*). Additionally, many pathways in metabolism are known to be controlled by signals from other compounds or nutrients. In bacteria, nitrogen, sulfur, phosphorus, and iron metabolism can even be controlled by carbon metabolism (*51*, *53*).

Our previous analysis of 332 yeasts identified a pervasive pattern of trait loss (*22*), which suggests that generalists have either retained carbon traits over a long evolutionary timescale or gained traits unlike their non-generalist relatives. To test these hypotheses, we used BayesTraits to compare the relative rates of carbon trait gain or loss either across all yeasts or specifically within a generalist background (Fig. 3D, table S7). For the eight carbon traits found in less than 75% of generalists, we identified strong trait loss across the entire phylogeny but some evidence of trait gain in the generalist background. Carbon generalists, therefore, appear to be able to both gain and retain carbon traits that are otherwise lost broadly across the rest of the subphylum.

Under the trade-off paradigm, carbon generalism would come at a metabolic cost such that specialists would grow faster on fewer carbon sources than the generalists, which would grow slower on more carbon sources. In contrast, our results revealed that carbon generalists were generally fast growers, nitrogen generalists, and able to retain and gain traits typically lost in non-generalists. Given the extreme carbon breadths of generalists and specialists, we hypothesized that these two groups would also differ in other aspects of metabolism, genome content, and ecology.

### Intrinsic factors drive generalism and specialism

Metabolic networks, including the carbon metabolism network, are highly interconnected due to shared enzymes and pathways. To examine whether metabolic network structure varied between generalists and specialists, we used Kyoto Encyclopedia of Genes and Genomes (KEGG) orthologs (KOs) to build metabolic networks for all yeast species and tested for a correlation between carbon breadth and six KEGG network features (Fig. 4A and fig. S6, table S8). Carbon generalists had a higher edge-count (i.e., more connections between nodes of the network) than carbon specialists (Fig. 4A) (*54*). Both carbon generalists and specialists had disassortative networks, a property of all biological networks (*55*), which reflected high levels of connection between nodes with dissimilar properties. However, the generalist networks were less disassortative (i.e., they had more highly interconnected nodes) than specialists (Fig. 4A). There were no significant correlations between carbon breadth and network modularity, diameter, betweenness, or density (fig. S6). Despite the extreme difference in carbon metabolism capabilities, carbon generalists and specialists had only slight differences in the size and shape of their global KEGG metabolic networks. These results suggest that generalist and specialist networks are overall similar in size and shape but differ in how they are wired.

**Figure 4:**
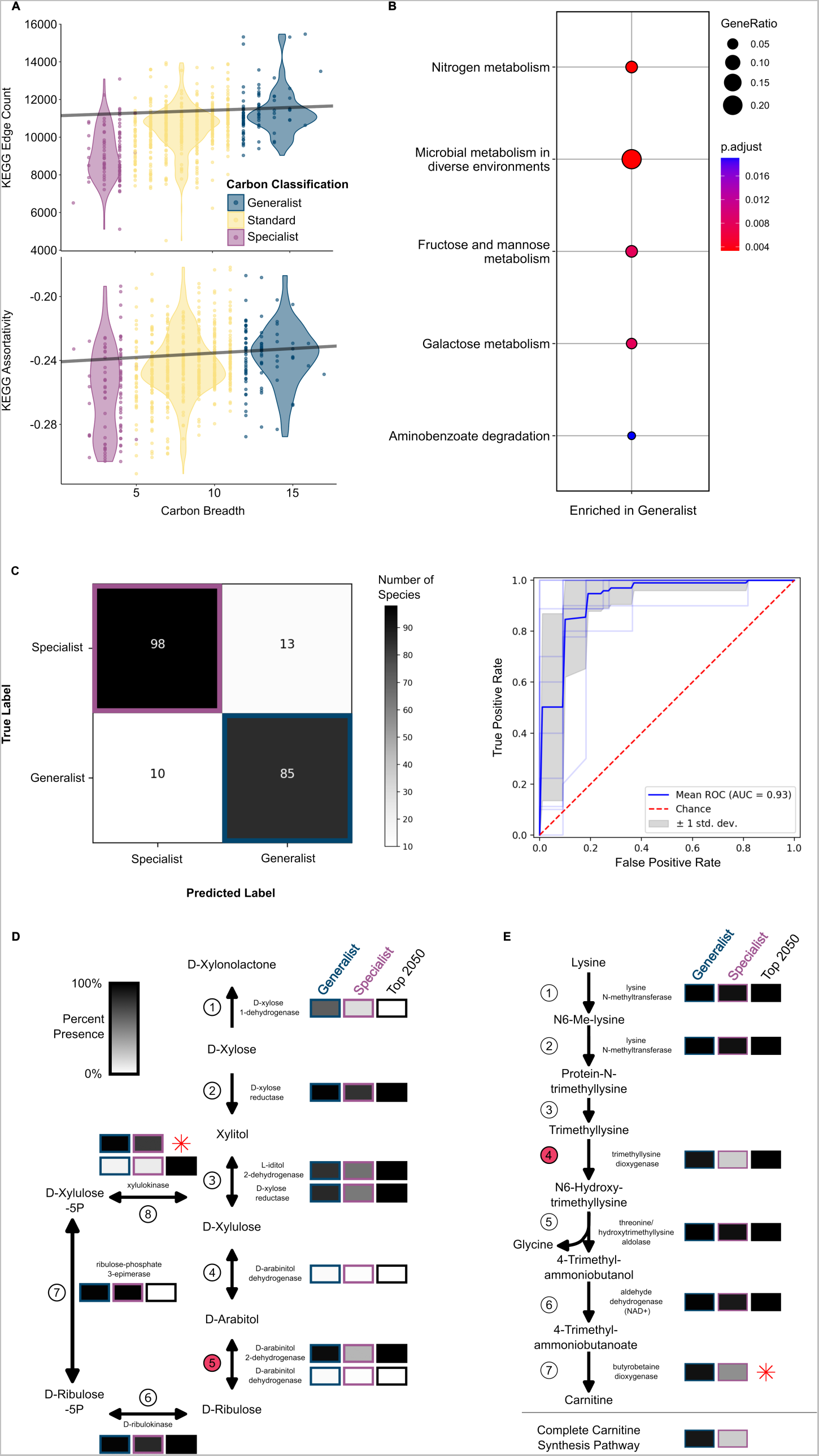
Generalist and specialist metabolism differs in expected and unexpected ways. A. Two KEGG network statistics were significantly and positively correlated with carbon breadth when taking into account phylogenetic relatedness (using phylogenetic generalist least squares regression). KEGG Edge Count (p-value 0.00014, slope 29.12) and KEGG Assortativity (p-value 0.00761, slope 0.000556) were both elevated in carbon generalists. B. KEGG enrichment analysis of KOs enriched in generalists (greater than 80% of individuals) and depleted (missing in < 20% of individuals) in specialists. The size of the circles represents the number of KOs per term relative to the number of KOs for that term. The color of the circle represents the FDR-corrected p-value. The analysis was compared to a background of all KOs present in generalist and specialist yeasts. C. The yeasts were classified into generalists and specialists using a machine learning algorithm trained on the KOs. The correct classification occurred in 88% of specialists and 89% of generalists. The ROC analysis suggested that both the sensitivity and specificity of our model is excellent-inln(AUC=0.93). D. Multiple reactions in the pentose and glucuronate interconversions pathway were important in classifying species into generalists and specialists (black boxes.) Boxes are shaded as the percent of each carbon classification with at least one enzyme in that step of the reaction. The reaction with the 3^rd^ highest relative importance in the machine learning analysis is shown in Step 5 and is facilitated by D-arabinitol 2-dehydrogenase. Interestingly, experimental studies suggest that this D-arabinitol 2-dehydrogenase in yeasts is capable of also completing the reaction in Step 4 (*114*). Step 8 was in the top features used in the machine learning analysis, despite the fact that KEGG only partially annotated this gene. The xylulokinase encoded in yeasts, *XYL3*, is well studied (*57*). Therefore, we re-annotated the *XYL3* gene and have shown the relative abundance of this gene (red star). E. Another pathway with multiple reactions that are important for classifying carbon generalists and specialists is the carnitine biosynthesis pathway. The reaction in Step 4 has the 4^th^ highest relative importance in the machine learning classification of carbon classification. Step 7 was not annotated by KEGG in any of our yeast species, but this step had been previously characterized in *Candida albicans* as being facilitated by the trimethyllysine dioxygenase enzyme encoded by *BBH2* (*62*). We re-annotated *BBH2* in our yeasts using this reference sequence and calculated the relative abundance in each carbon classification (red star). Finally, we determined the number of species that could hypothetically complete the lysine to carnitine biosynthesis pathway.

We next investigated differences in the composition of generalist and specialist networks. Generalists and specialists largely showed similar compositions across KOs, but a small set of KOs were depleted (presence < 20%) in specialists and enriched (presence >85%) in generalists (Fig. 4B, table S9). Generalist-enriched KOs were related to nitrogen, fructose, mannose, and galactose metabolisms. Enrichment of these terms suggests that differences in gene content contribute to the overall differences observed between generalists and specialists for carbon metabolism traits.

To further understand the relationship between niche breadth and gene content, we employed machine learning. Specifically, we trained a supervised random forest classifier to use KO presence and absence as predictive features for niche breadth classification. The resulting classifier was both highly sensitive and specific, correctly classifying 88% of specialists and 89% of generalists (AUC=0.93; Fig. 4C). The high accuracy suggests that generalist and specialist KEGG networks differ in ways that were not detected in the KO enrichment analysis.

Examination of features that the classifier relied on to accurately predict specialists and generalists using dropout analysis identified 2,050 KOs that significantly contributed to classification accuracy. Approximately 5,000 unique yeast KOs were used to train the algorithm, suggesting that many KOs contributed some information to niche breadth classification. We further examined the top four features because the fifth feature had only half the relative importance score of the other four. Two of the top four features had direct links to the catabolism of specific carbon substrates, demonstrating the power and precision of our algorithm. The KO for *manB (*K01192), which encodes a *β*-mannosidase, had the second highest relative importance (relative importance 0.048). This KO was identified in 7% of specialists (8/111) and 80% of generalists (76/95). *β*-mannosidases are known to have a role in microbial utilization of *N*-glycans as a carbon source (*56*). Almost all the carbon generalists (93/95) can utilize mannose, which leads to the hypothesis that generalists likely use the mannose moieties present in *N*-glycans as a carbon and energy source.

The KO with the third highest importance was K17738 (relative importance 0.043), which is the *ARD* gene encoding D-arabinitol 2-dehydrogenase, an important component of the pentose and glucuronate interconversions pathway (Fig. 4D, step 5). This KO was more frequently present in the genomes of generalists (96%, 91/95) than in the genomes of specialists (71%, 79/111). Indeed, we found a portion of that pathway where 5 of the 8 reactions were among the 2,050 KOs (with two falling in the top 100 KOs) that contributed to the classification of carbon generalists and specialists (black boxes in Fig. 4D). Importantly, growth on xylose was included in our carbon classification, and the xylose metabolism genes *XYL1* (Step 2 in Fig. 4D), *XYL2* (Step 3), and *XYL3* (Step 8) were all identified as important features (with *XYL1* falling within the top 100), suggesting that xylose metabolism genes are promiscuous and may have multiple metabolic capabilities (*57*). This result also supports the hypothesis that intrinsic genetic factors contribute to niche breadth by connecting pathways.

The feature with the highest relative importance was K03940 (relative importance 0.062), which encodes an NADH ubiquinone oxidoreductase core subunit (NDUFS7 in humans) of Complex I in the mitochondrial electron transport chain. This KO was identified in 29% of specialists (32/111) and 95% of generalists (90/95). Interestingly, Complex I is known to vary widely, in presence and makeup, including the presence of an alternative pathway in some yeasts. In *S. cerevisiae*, the NADH oxidoreductase function of Complex I is conducted by three single-subunit enzymes (Ndi1p, Nde1p, or Nde2p) (*58*). Conversely, in *Y. lipolytica,* Complex I is composed of 42 subunits, including the NADH ubiquinone oxidoreductase NUKM (K03940) (*59*). Thirty additional Complex I enzymes were within the top 2,050 KOs, and two fell within the top 10%: K03941 and K03966, which are both NADH ubiquinone oxidoreductases in the *β* subcomplex (KEGG Map map00190). While the complete loss of canonical Complex I in the Saccharomycetales, an order with many specialist yeasts, contributes to the high relative importance of the K03940, the effect is seen in other orders. For example, within the Pichiales, 100% (5/5) of generalist genomes encode K03940, in contrast to only 18% (6/33) specialists. Complex I has been implicated in *C. albicans* growth and virulence (*60*), as a global regulator of fungal secondary metabolism in *Aspergillus* (*61*), and results in a higher proton motive force compared to the alternative pathway in *S. cerevisiae*. The presence of Complex I in generalists, therefore, may support increased carbon breadth and elevated growth rates.

The last KO we investigated was K00474 (relative importance 0.043), which is a trimethyllysine dioxygenase involved in lysine degradation. Every step in the pathway that degrades lysine to carnitine, except the last step, was identified as important in the machine learning classification. The last step (Fig. 4E, Step 7) was not annotated by KEGG in any of our yeasts. Therefore, we annotated the *BBH2* gene, which encodes the trimethyllysine dioxygenase, directly from our predicted coding sequences using previously published reference sequences (Figshare repository) (*62*). After manual annotation of *BBH2*, we found that most carbon generalists (*63*) were predicted to be able to complete the carnitine biosynthesis pathway (91%: 86/95), while very few carbon specialists were predicted to do so (20%: 22/111). Carnitine plays an important role in the transport of acetyl coenzyme A (acetyl-CoA), which in turn is a major metabolite that contributes to many metabolic pathways, including the production of ATP in the mitochondrial tricarboxylic acid (TCA) cycle. Acetyl-CoA can be supplied to the mitochondria through direct production within the mitochondria when glucose is available or, when glucose is unavailable, it can be transported into the mitochondria using the carnitine shuttle (*63*). Some yeasts, including *C. albicans,* rely solely on the carnitine shuttle for this transport (*62*), while other yeasts, such as *S. cerevisiae,* can utilize a carnitine-independent method for acetyl-CoA transport (*64*). Similarly, some yeasts, such as *C. albicans*, can synthesize carnitine; others, such as *S. cerevisiae*, cannot and rely on exogenous sources. A complete carnitine synthesis pathway may ensure acetyl-CoA transport when glucose is unavailable, especially in species that rely solely on the carnitine shuttle.

Additionally, carnitine and the carnitine acetyltransferases can be essential for growth on some nonfermentable carbon sources. These include ethanol in all cases, as well as glycerol in certain mutants with disrupted citrate metabolism (*65*). We found that 90.5% (86/95) of generalists can grow on glycerol compared to only 24.5% (27/110) of specialists (table S2). Moreover, specialists that can grow on glycerol are more likely to have the complete carnitine synthesis pathway than those that do not (z-test, χ^2^= 10.425, p-value = 0.0186). These results suggest that carnitine production is another important trait that affords metabolic flexibility and carbon breadth.

### Ecological overlap between generalists and specialists

In addition to intrinsic factors, extrinsic factors could also shape yeast metabolic niche breadth. If habitat were a major driving force in the evolution of generalists and specialists, we would expect the two carbon classifications to partition by environment. To test this hypothesis, we retrieved environmental data for 1,088 yeast strains using a novel formal ontology. We analyzed environmental isolation data for 743 yeasts with both isolation and growth classification data. We analyzed either all direct classes of isolation environments or one of five mutually exclusive high-level environments: Arthropoda (210), Chordata (50), Environmental isolates (such as soil, water etc.: 149 yeasts), Plant (268), and Victuals (defined as processed food or drink: 66 yeasts).

At all levels of our ontology, generalists and specialists shared isolation environments. For example, *Hyphopichia homilentoma* (generalist) and *Wickerhamomyces sydowiorum* (specialist) were both isolated from the tunnel of the wood-boring beetle *Sinoxylon ruficorne* in the red bushwillow *Combretum apiculatus*. Some yeast pairs were isolated from the same species of plant but different parts of the tree, such as *Sugiyamaella americana* (generalist) from the oak tree *Quercus kelloggii* and *Kregervanrija fluxuum* (specialist) from the sap of *Q. kelloggii*. As a final example, *Candida bentonensis* (generalist) was isolated from apple cider, while *Pichia nakasei* (specialist) was isolated from apple must (apple cider with the skin).

Analysis of the full ontology of isolation environments also found generalists and specialists in the same broad isolation environments. We built a random forest algorithm to classify carbon classifications when trained on the isolation environment ontological data (fig. S7), but it was only slightly better than chance (AUC = 0.57) and correctly classified 59% of specialists (59/100) and 52% of generalists (47/91). Moreover, we did not find significant differences in the numbers of generalists and specialists across the high-level environmental classifications (Fig. 5A). This result suggests that, either environment does not play a major role in shaping carbon niche breadth, or habitat partitioning is occurring within a shared environment by competitors consuming different carbon sources within that environment.

**Figure 5:**
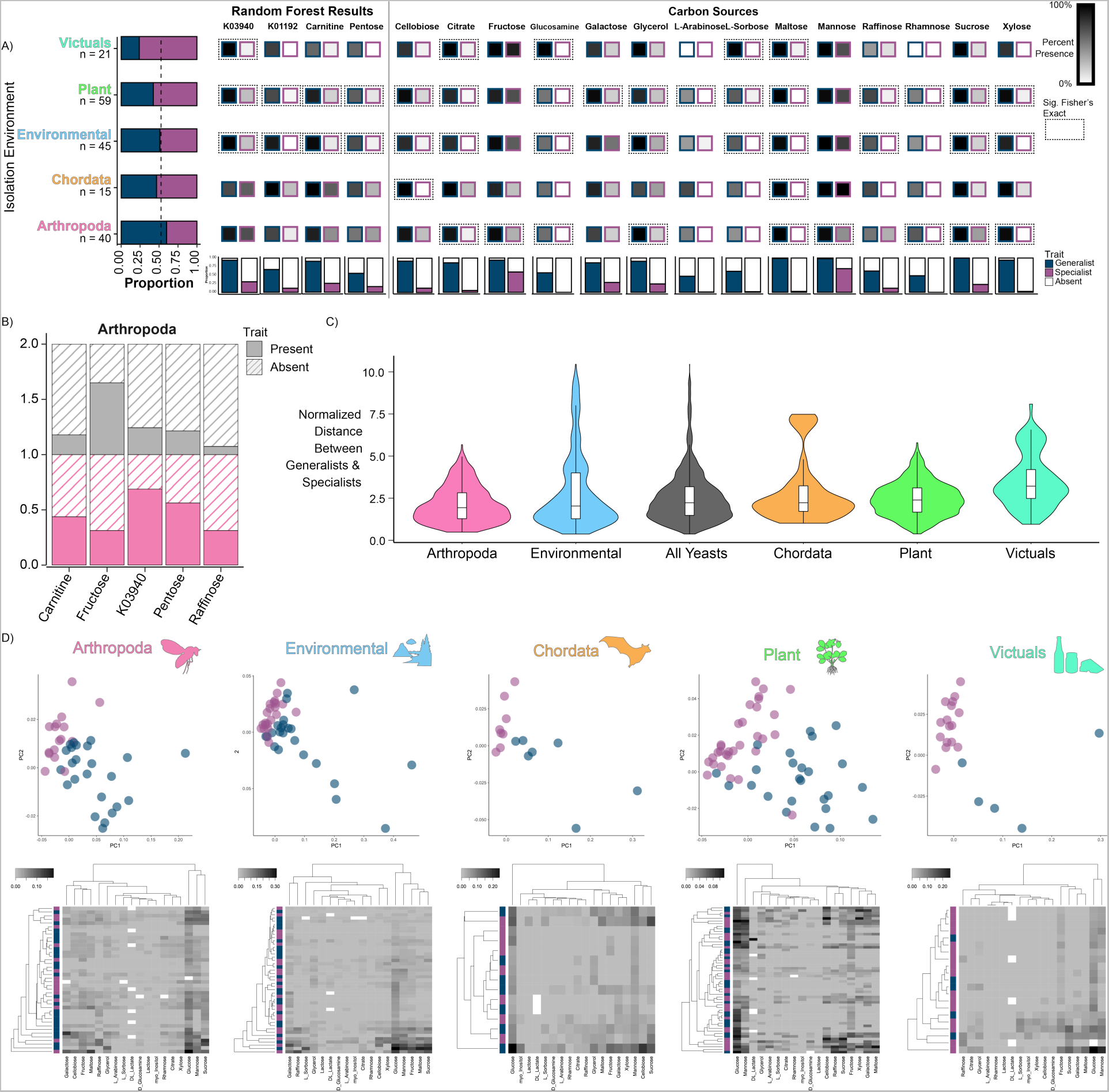
Isolation environments host both generalists and specialists. A. There was no significant difference in the number of generalists and specialists isolated from the five major isolation environmental classes. The bar graph on the left-hand side shows the proportion of generalists and specialists isolated from each environment. The dashed line represents the 50% line. B. Specialists show variation in trait presence based on isolation environment. Bar graph comparing trait presence (solid color) of specialists in the Arthropoda environment (pink) relative to all other environments (gray). All results shown are significant. C. Generalists and specialists were most similar with respect to growth rate when isolated from Arthropoda and least similar when isolated from Victuals. The violin plots visualize the normalized distance between generalists and specialists found in a specific environmental class in the first 4 principal components of a phylogenetic principal components analysis (pPCA, cumulative variance 78%) constructed from continuous growth rate data on 18 carbon substrates. D. The heatmaps show the growth rates of species isolated from each environment with the species clustered on the y-axis by their distances in the pPCA and the carbon sources clustered on the x-axis by growth rates. Missing data are shown in white.

To test whether generalists and specialists were metabolically partitioning within environments, we used a Fisher’s Exact test to compare species counts for the ability to grow on different carbon and nitrogen sources within an environment. We found that, for most traits, generalist and specialist either did not differ in their metabolic capabilities or showed differences driven by the generalist phenotype (Fig. 5A). We next quantified differences in growth within generalists and specialists for traits across environments. We found that, within specialists, the ability to grow on fructose was depleted in the Arthropoda environment (p = 0.046, Fig. 5B) but enriched in the Victuals environment (p = 0.016, Fig. 5B). These results suggest that, within the Arthropoda environment, generalists and specialists could be partitioning their niches differently. Of course, we cannot rule out the possibility that generalists and specialists partition niches at finer scales than those at which isolation data were collected.

Although generalists and specialists did not generally differ in their ability to grow on carbon sources within similar environments, we hypothesized that there might be differences in growth rates. To test this hypothesis, we performed phylogenetically corrected principal components analyses (pPCAs; fig. S8) based on carbon source growth rates. We then measured and normalized the distance between generalists and specialists. Generalists and specialists isolated from Arthropoda were more similar to each other than generalists and specialists isolated from all other environments (Wilcoxon rank sum, p-value= 1.59×10^−14^.) This result is reflected in the nearness of points in the pPCAs and in the heatmaps (Fig. 5D). In contrast, in the Plants and Victuals environments, generalists and specialists were more different from each other than generalists and specialists isolated from all other environments (Wilcoxon rank sum, Plants p-value= 0.00066, Victuals p-value= 2.20×10^−16^.) Interestingly, in the Victuals, generalists and specialists were non-overlapping in the pPCA (Fig. 5D).

Our results suggest that carbon generalism and specialism are both viable strategies for adaptation to a wide range of habitats. The Arthropoda environment may be shaping generalists and specialists in similar ways as generalists and specialists were more similar than expected in this environment. Conversely, the Victuals environment included relatively few generalists, and those generalists differed greatly from the specialists isolated from Victuals, which likely reflects strong anthropogenic influence.

### The power of the Y1000+ Project dataset

We developed a rich ensemble of genomic, metabolic, and environmental data that, when coupled with a powerful analytical framework, enabled us to understand the evolution of ecological niche breadth across an entire eukaryotic subphylum. This comprehensive dataset and state-of-the-art analytical framework provide the opportunity to study numerous other complex traits. To illustrate this potential, we examined how yeast metabolic niche breadth intersects with human pathogenicity. The World Health Organization (WHO) recently released its first-ever fungal priority pathogens list (FPPL), which included six Saccharomycotina species (*66*). We defined 11 yeasts as opportunistic human pathogens because they are known to cause human infections and generally require biosafety level 2 (BSL-2) precautions in research laboratories. For comparison, we also identified close relatives of pathogenic species using a specific phylogenetic distance cutoff to standardize the clades. We then investigated differences between pathogenic yeasts and their relatives using Y1000+ Project data (Fig. 6).

**Figure 6:**
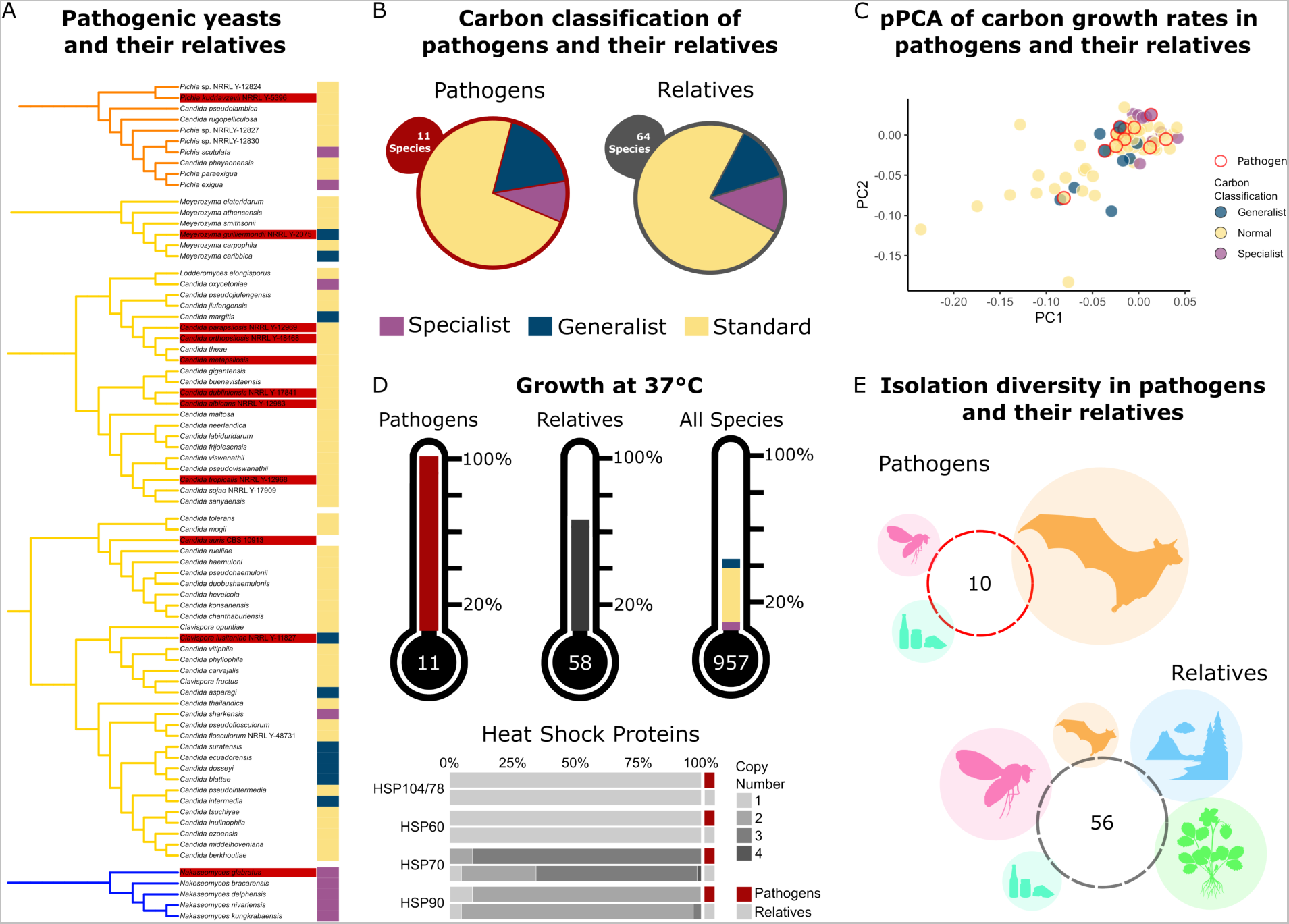
Carbon generalism and specialism is not associated with pathogenicity in yeasts. **A.** The phylogenetic clades containing human fungal pathogens. Clades reflect all species within a specific phylogenetic distance from the identified pathogen. Pathogens are found in 3 different orders, and at least one is generalist, specialist, and standard. **B.** Pathogens and their relatives had nearly identical proportions of generalist, specialist, and standard yeasts. This result suggests that carbon breadth is not a defining or predictive factor for the potential of a species to gain the ability to infect humans. **C.** Pathogens and their relatives did not differ substantially in their growth rates on carbon substrates. The pPCA was constructed using growth rates on carbon substrates and projected onto the first two components (totaling 80**%** of the total variance.) Pathogens did not cluster together, while generalists and specialists appeared further apart. This result suggests that pathogens do not have shared growth rate characteristics. **D.** Proportion of yeasts that can grow at 37°C in pathogens, their relatives, and all sampled yeasts. All yeasts identified as pathogens can grow at 37°C. Pathogenic yeasts as compared to their relatives have a statistically different proportion of yeasts that can grow at 37°C (Chi-Squared Test, p = 0.042). Heat shock protein (*HSP*) gene copy number was determined using Interpro and KOs. *HSP* gene copy number was not associated with pathogenicity. **E.** Isolation environment for the specific strains of pathogens and their relatives. Circles are proportional to the percent of species isolated from Chordata (orange), Arthropoda (pink), Victuals (teal), Environmental (blue), and Plants (green.)

Carbon sources and availability vary *in vivo* in humans, suggesting that carbon breadth may play an important role in promoting or preventing fungal pathogenesis (*67*). The proportion of pathogenic yeasts classified as standard, generalist, and specialist yeasts was similar to that of their non-pathogenic relatives (Fig. 6A). We also observed extreme carbon breadths within pathogenic yeasts: *Meyerozyma guilliermondii* had a carbon breadth of 15, while *Nakaseomyces glabratus* (syn. *Candida glabrata*) (*68*) had a carbon breadth of 2. This result supports previous findings that, while carbon metabolism is vital for fungal pathogens as they assimilate to a host, the mechanisms may vary widely, especially in *N. glabratus* (*69*).

Previous work in *C. albicans* linked its pathogenicity to its high growth rate (*70*). To examine this idea, we visualized the pathogenic yeasts and their relatives on a pPCA using all our collected growth rate data (Fig. 6C). We observed no clustering of pathogenic yeasts using carbon growth rates. Moreover, fungal pathogens within the same relative group varied in their growth rate on glucose by almost 3-fold: *Candida parapsilosis* had a growth rate of 0.042, while *Candida tropicalis* had a growth rate of 0.124.

Our growth rate data, however, were collected at room temperature in a defined medium and may not reflect growth rates in human infections. One feature known to be necessary, but insufficient, for pathogenicity is growth at human body temperature or 37°C (Fig. 6D) (*69*). We also observed that relatives of human pathogens had an elevated rate of growth at 37°C (~64%) compared to all species for which growth at this temperature was measured (~41%). This result likely reflects the necessity of growth at 37°C to evolve prior to pathogenicity.

Genomic differences between pathogenic yeasts and their relatives can also be investigated. It was previously shown that heat-shock proteins (HSPs) are likely involved in growth at high temperatures in *C. albicans* (*71*). Examination of copy number variation in the genes encoding HSPs in the pathogenic species and their relatives (*71*) identified an increase, albeit slight, in *HSP70* gene copy number among pathogenic yeasts (Fig. 6D).

The non-human environments in which we find opportunistic fungal pathogens may select for traits that promote opportunistic pathogenicity (*72*). Moreover, the ecology of fungal pathogens provides insight into exposure risk of exogenous infection from natural reservoirs. While many fungal infections are nosocomial (acquired in a health-care facility) or arise from commensal associations, pathogenic yeasts can also be isolated from the environment (*73*, *74*). Nonetheless, we know little about their ecologies outside of clinical settings (*73*, *74*). Eight of the 11 fungal pathogen strains examined here were isolated from humans, one from insect frass, and one from fruit juice, while the origin of *Candida orthopsilosis* is unknown (Fig. 6E). Relatives of the pathogenic species were isolated from all our environmental classifications, including five that were isolated from human specimens. The lack of notable differences between yeast pathogens and their non-pathogenic relatives mirrors findings from other clades and provides support for the hypothesis that the traits and genetic elements contributing to pathogenicity are not broadly shared across pathogens but unique to each (*75*). Fungal pathogens are largely opportunistic, suggesting that many of the traits and genetic elements that contribute to their pathogenicity likely evolved for survival in natural environments and not inside human hosts (*75*). Therefore, understanding the evolution of fungal pathogenicity requires understanding of the natural history, ecology, and adaptations of fungi that facilitate their success in their natural environments. Although understanding how ecology contributes to the evolution of fungal pathogens will require substantial additional research, the rich empirical and analytical framework presented here shows considerable potential for the study of fungal pathogenicity and other complex traits on a macroevolutionary scale.

## Conclusions

We generated an unprecedented dataset of genomic, metabolic, and ecological data for nearly all known species of the 400-million-year-old yeast subphylum Saccharomycotina (*22*, *48*). This dataset increases the number of species with available genomes for the subphylum by over 3-fold (making it the first eukaryotic subphylum with near-complete taxon sampling), provides quantitative growth data for 853 strains across carbon and nitrogen conditions, and lays out a novel ecological ontology for 1,088 yeasts that is portable to other microbes and beyond. In addition to familiar yeasts, such as *S. cerevisiae* and *C. albicans,* this dataset highlights 1,049 species that display vast genomic, metabolic, and ecological diversity. We examined the genomic, phenotypic, and ecological variation found within this fungal subphylum using a powerful analytical framework of machine learning, phylogenomic, and network approaches to address several outstanding fundamental questions about how generalists and specialists function and evolve. We found that generalists typically grew faster on carbon sources than specialists. The carbon metabolism traits found within generalists were either maintained across evolutionary time or gained, even though there was a strong overall trend for trait loss across the subphylum. Despite extreme variation in carbon metabolic capabilities, we found no large-scale metabolic differences between generalists and specialists. Machine learning allowed us to identify specific genes and pathways shared by generalists but largely absent from specialists. These pathways or complexes are directly involved in carbon and energy metabolism, often by enhancing metabolic flexibility and robustness, a key result that ties ecological variation to genomic variation. Finally, we found that both generalists and specialists were isolated from the same environments. These results suggest that generalism and specialism are driven mainly by intrinsic factors, rather than extrinsic factors or trade-offs. We propose that it is the accumulation of genetic (intrinsic) changes that gives rise to the large-scale phenotypic differences of generalism and specialism.

Here we focused on answering questions about the evolution of generalism and specialism, but the potential for this dataset is vast. For example, we also highlighted how other complex traits, such as pathogenicity, could be studied using the dataset and a similar analytical framework to the one we used in this study. Aside from studying the evolution of specific complex traits, additional questions that could be addressed with these data include quantifying correlations among genes, traits, and/or ecologies; investigations of gene family evolution; and genome-informed bioprospecting of yeasts and their genes for the sustainable production of cellulosic biofuels and bioproducts. By coupling a comprehensive dataset with a robust analytical framework, the Y1000+ Project provides a roadmap that connects DNA to diversity.

## Materials and Methods

### Strain Information

Taxonomic type strains of yeast species were obtained from the USDA Agricultural Research Service (ARS) NRRL culture collection in Peoria, Illinois, USA (n = 777) and the Westerdijk Fungal Biodiversity Institute CBS culture collection in the Netherlands (n = 250); a small number of yeasts came from other labs (table S1A). Cultures were sent from the culture collections as frozen stocks in glycerol yeast extract peptone dextrose (YPD), on YPD slants, or as lyophilized cultures that were revived in YPD upon arrival and cultured onto YPD plates for single colonies to be grown to saturation in YPD to be frozen down in 15% glycerol at −80°C. For 55 yeast species where we could not obtain the taxonomic type strain, we used instead another available strain with a preference toward authentic or widely used reference strains (table S1A). In addition to the named yeast species, 62 taxa included in our dataset are candidates for novel species that await formal taxonomic description; 28 were isolated in the Hittinger Lab at the University of Wisconsin – Madison using standard protocols (*76*, *77*), while the remaining 34 were obtained from the ARS NRRL Culture Collection. Species names were up-to date as of May 2023. For synonymous names and recent taxonomic updates, we refer the reader to the online MycoBank database. It is important to note that, for all analyses, a single strain of most species was used. There are many strains of species available in culture collections. The use of a single strain means we have not fully explored species- or population-level genomic, phenotypic, or ecological variation.

### Genome Sequencing and Assembly

For approximately half the species, genomic DNA was extracted and fragmented using a phenol:chloroform extraction protocol, followed by sonication (*22*, *78*). Libraries were prepared using the NEBNext Ultra II DNA Library Kit using Illumina adaptors. For the rest, the protocol was slightly modified to remove the sonication step because an enzymatic fragmentation step was used during the library prep using the NEBNext Ultra II FS DNA Library Kit. All paired-end libraries underwent paired-end sequencing (2×250 base pairs) on an Illumina HiSeq 2500 instrument.

Paired-end Illumina DNA sequence reads were used to generate whole genome assemblies using the meta-assembler pipeline iWGS v1.1 (*79*). The raw sequenced reads were processed by trimming adapters and low-quality bases with Trimmomatic v0.33 (*80*) and Lighter v1.1.1 (*81*). Next, the optimal k-mer length for each genome’s assembly was determined by KmerGenie v1.6982 (*82*). We used the processed sequence reads as input into six different de novo assembly tools: ABySS v1.9.0 (*83*), DISCOVAR r51885 (https://software.broadinstitute.org/software/discovar/blog), MaSuRCA v2.3.2 (*84*), SPAdes v3.7.0 (*85*), and Velvet 1.2.10 (*86*). The genomes of yeast species varied in their ploidy and levels of heterozygosity. We identified genome assemblies that required ploidy-aware assembly by manually examining the distribution of variable site frequencies. Distributions of variable site frequencies were calculated using BAM-formatted files as input to nQuire, version a990a88 (*87*). Prior to examining variable site distributions, denoising was conducted to minimize the impact of erroneous read mapping. For genomes with distributions of variable sites that had two or more peaks (suggestive of containing two or more sets of chromosomes and/or high levels of heterozygosity), the original assemblies were further processed by HaploMerger2 v20180613 (*88*) to collapse allelic contigs and obtain consensus assemblies. Genomes processed with HaploMerger2 are labeled with the suffix “haplomerger2” in the genome assembly names.

The quality, genome size, and GC content of all genome assemblies was assessed with QUAST v5.0.2 (**table S1A**) (*89*). The ‘‘best’’ assembly for each genome was identified as the one with the highest N50 value and an appropriate genome size. We evaluated the completeness of all 1,154 genome assemblies using the Benchmarking Universal Single-Copy Orthologs (BUSCO), version 5.23.2 0 (*90*). Each assembly’s completeness was assessed based on the presence and absence of a set of 2,137 predefined orthologs that are single-copy in at least 90% of the 76 reference yeast genomes in OrthoDB v10 (*91*, *92*). The BUSCO statistics revealed an artifactual level of duplication in some DISCOVER assemblies, so we replaced them with SPADES assemblies and marked their assembly names accordingly.

#### Genome Filtering

Mitochondrial DNA contigs were removed from each nuclear genome assembly based on homology to reference yeast mitochondrial sequences and the presence of BUSCO gene annotations. Briefly, we identified putative mitochondrial contigs based on blastn hits against both the complete reference mitochondrial genomes (coverage >30%) or their coding sequences (coverage >70%). Putative mitochondrial contigs that contained no nuclear BUSCO annotations were removed.

To examine if the genomes had significant levels of bacterial contamination, we compared the genomes to the NCBI reference prokaryotic database (ref_prok_rep_genomes.00). We then used blastn with an e-value cutoff of 1e-11 to compare the reference bacterial genomes to the assembled fungal genomes. Fungal contigs were identified as potential contaminants if they had over 50% coverage and over 50% percent identity to a bacterial sequence. Nine genomes were identified as having sufficient contamination to warrant additional filtering. The contaminated contigs were removed from subsequent analyses. All genomes were also filtered of contigs less than 500bp.

The filtered genomes can be found on the Figshare repository. Each genome was also uploaded to GenBank, which utilizes an independent filtering algorithm. All analyses conducted in this manuscript were done on the genomes housed on the Figshare repository.

### Genome Annotation

For each of the 1,154 yeast genome assemblies (including previously sequenced genomes), repetitive sequences were identified and softmasked using RepeatMasker v4.1.2 with the “-species” option set to “saccharomycotina”. Protein-coding genes were annotated using BRAKER v2.1.6 (*93*) using all Saccharomycetes protein sequences in the OrthoDB v10 as homology evidence and the ab initio gene predictors AUGUSTUS v3.4.0 (*94*) and GeneMark-EP+ v4.6.1 (*95*). The BRAKER pipeline was run in the EP mode to process all protein homology-evidence using ProtHint v2.6.0 (*95*), and the “--fungus” option was turned on to run GeneMark-EP+ with the branch point model for fungal genomes. For genes with multiple transcripts, only the longest transcript was retained. In yeasts, the CUG codon can encode the canonical leucine or one of two alternatives, serine or alanine. In clades with alternative nuclear genetic codes, CUG is rare, so the peptide translations were all conducted using the standard nuclear codon table (Genetic Code Translation Table 1).

Gene models predicted using the BRAKER pipeline were functionally annotated using the KEGG (*54*) and InterPro (*96*) databases. KEGG orthologs (KOs) were assigned to protein sequences of gene models using the KofamScan tool (*97*) run in mapper mode (“-f mapper”) and without reporting unannotated sequences (“--no-report-unannotated”). InterPro annotations were assigned to protein sequences of gene models using the InterProScan 5.52-86.0 tool (*98*) run with all the available analysis modules, as well as the lookup of GO terms (“--goterms) and Pathway annotations (“--pathway). Every protein sequence was annotated locally, to avoid any potential bias in the lookup of precalculated annotations (“–disable-precalc”).

While the standard genetic code was used to translate the protein annotations, there are three independent nuclear codon reassignments within the subphylum. To validate previous findings and identify any possible novel nuclear genetic codes, we used the Codetta method (*99*). Analyses were run using the standard parameters. Nuclear genomic tRNA contents were estimated using tRNAscan-SE 2.0.9 using the standard eukaryotic parameters.

#### Genome Analyses

We previously identified 2,408 orthologous groups (OGs) of genes that were nearly universal among 332 yeast species representing all 12 orders (*22*). To retrieve orthologs of these 2,408 molecular markers from all 1,175 proteomes (1,154 yeasts and 21 selected outgroup taxa), we used orthofisher v1.02 (*100*). To minimize the inclusion of potentially spurious or paralogous genes for each of the 2,408 OGs, we reconstructed a fast maximum likelihood (ML) tree using the program FastTree version 2.1.9 (*101*) with the LG model of amino acid substitutions (*102*). We then discarded all sequences whose terminal branch (leaf) lengths were at least 20 times longer than the median of all terminal branch lengths across the fast ML tree for each OG.

To construct the phylogenomic data matrix, we first aligned the amino acid sequences of each of the 2,408 OGs using MAFFT v7.299b (*103*) with the options ‘‘--thread 8 --auto --maxiterate 1000’’. Next, we trimmed alignments using the trimAl v1.4.rev15 (*104*) with the options “-gappyout”. Lastly, we retained only those OGs that had ≥ 50% taxon occupancy and alignment length ≥ 200 amino acid sites, which left a total of 1,403 OGs (719,591 amino acid sites).

We inferred the concatenation-based ML tree using IQ-TREE v2.0.7 (*105*) on a single node with 100 logical cores under a single “LG +G4” model with the options “-T 100 -mem 800G -m LG+G4 -bb 1000”. We chose the “LG +G4” model because 726 / 1,403 genes favored it as the best-fitting model. We ran 5 independent tree searches to obtain the best-scoring ML gene tree. We also inferred the coalescent-based species phylogeny with ASTRAL-III version 4.10.2 using the set of 1,403 individual ML gene trees inferred by IQ-TREE. The reliability of each internal branch was evaluated using 1,000 ultrafast bootstrap replicates and local posterior probability, in the concatenation- and coalescence-based species trees, respectively. A time tree was estimated using the RelTime method, as implemented in the command line version of MEGA7 (*106*). Calibration points were adopted from our previous work (*22*). The time tree and node times are available on the Figshare repository. We visualized and plotted phylogenetic trees using iTOL v5 (*107*). Multiple sequence alignments and phylogenetic trees are available on the Figshare repository.

### Phenotyping

To determine whether there was metabolic variation across yeast species or clades, we generated a quantitative growth dataset for 853 yeasts across 24 different carbon and nitrogen substrates. We tested growth for all yeast species in our dataset, except for the plant pathogens in the genus *Eremothecium*. However, some yeasts did not produce reliable growth data. A subset of yeasts did not grow or had delayed growth that went beyond the length of our growth experiments. Among the yeasts that did grow, some had noisy growth data, likely due to flocculation or some other artifact of growth, and their growth rates could not be quantified. To generate growth data for carbon and nitrogen traits, we measured growth phenotypes of yeasts across eight sets of growth experiments in triplicate. Each set consisted of approximately 192 species. Yeasts were streaked for single colonies on yeast extract peptone dextrose (YPD) plates. Each week, a single colony was picked and inoculated in YPD liquid medium and allowed to grow for one week on a spinning drum at room temperature to ensure the yeasts were saturated. For carbon growth assessments, all species were then inoculated into 96-well plates containing 1% carbon source plus ammonium sulfate minimal medium (6.7g/L yeast nitrogen base w/o carbon, amino acids, or ammonium sulfate; 5g/L ammonium sulfate). For nitrogen growth assessment, all species were similarly inoculated into 1% glucose minimal medium (6.7g/L yeast nitrogen base w/o carbon, amino acids, or ammonium sulfate; 1% glucose), while varying the nitrogen source; nitrogen sources were normalized for the amount of nitrogen atoms present in the compound (*44*). Fresh media were made for each replicate and set. After a week of growth at room temperature, the cells were inoculated into fresh media in a 96-well plate. We found that some yeast species can store nutrients, which allows them to grow even if they cannot use the predominant sugar available in the media. Therefore, we performed two rounds of growth in the carbon sources of interest to limit this effect; we also included a no carbon control to determine whether yeasts were still able to grow on residual nutrient stores. All plates were placed on a BMG Omega SpectroStar Plate at room temperature to measure the optical density for one week at room temperature (Figshare repository). Across each replicate, species and media were randomized across and within plates to limit plate edge effects due to evaporation. Additionally, to limit evaporation we used Thermo Scientific™ Nunc™ Edge™ 96-Well, Non-Treated, Flat-Bottom Microplates, which are moated plates. We placed sterile water around the perimeter of the wells to limit evaporation. We previously published a subset of this data for specific carbon sources and species (comprising <5% of the total dataset) in smaller-studies (*57*, *108*). Growth rates were calculated in R using the *grofit* (v. 1.1.1-1) package (*109*). All growth curves were visually assessed to determine whether the species grew on a carbon or nitrogen source. Additionally, each growth curve was compared to the no carbon control for that species to determine whether the yeast was using stored nutrients or using the carbon sources provided. Species were dropped from the growth experiments if they either had noisy growth curves due to flocculation, another growth artifact, or did not grow during the experiment. Most yeasts grew and saturated within the first 72 hours of growth. However, there were 23 yeast species that displayed delayed growth; these either saturated after growth for seven days or did not saturate during the growth experiment. Any carbon or nitrogen source where there was no growth was set to zero to remove noise and background readings from the plate reader, and averages of replicates were calculated. To determine whether yeasts preferred sugars other than glucose, the growth rate of a yeast species on specific sugars was divided by that species’ growth rate on glucose. For any sugar where the normalized growth rate was greater than one, the yeast was considered to potentially prefer that carbon source over glucose. Proportions of glucophily and non-glucophily were calculated in R. For some analyses, growth rates were converted to binary data (growth or no growth).

A subset of our growth experiments was independently validated in a second laboratory. Specifically, we retested 16 strains of yeasts that were found to be fructophilic in our original growth experiments for this capability. All experiments were completed using the methods described above in both 1% glucose and fructose minimal media with minor modifications. Growth experiments were performed at 23°C in a plate reader, which was read every 30 minutes for seven days. Results of this experiment were compared to the previous growth experiments to confirm that the yeasts preferred fructose relative to glucose.

Finally, binary *C. auris* data was added to the final analysis conducted on fungal pathogens based on information gathered from the CBS strain database (https://wi.knaw.nl/).

### Ecological Niche Breadth Classification and Evolution

To classify generalists and specialists, we first calculated niche breadth from our binary growth data for each species. We generated a breadth matrix and then generated 1,000 permuted matrices. The permuted matrices were generated in R using picante (v. 1.8.2) (*110*). Randomizations were performed using the *randomizeMatrix* function with the null model “frequency” method. The breadth for a species was calculated for each randomized matrix, and we calculated the binomial confidence intervals to determine whether there was a significant difference between the observed breadth for a species relative to the randomized breadth using the R package Hmisc (v. 4.7-1). Species that had an observed breadth significantly greater than the distribution of randomized breadths were classified as generalists (between 12-17 and 4-6 (plus ammonium) for carbon and nitrogen sources, respectively); species that had a significantly lower observed breadth were classified as specialists (between 1-5 and 0 (ammonium only) for carbon and nitrogen sources, respectively); the rest of species were classified as standard. There were 30 yeasts that can grow on five carbon sources, which were classified as standard for carbon growth because of insignificant p-values. All calculations and classifications for carbon and nitrogen generalists and specialists were done separately. All summary statistics for carbon and nitrogen classifications were calculated in R, and all graphs were made in ggplot (v. 3.3.6).

We used BayesTraits v4.4.0 (http://www.evolution.reading.ac.uk) to model carbon and nitrogen breadth across the 852 species with measured breadths; one species was removed due to branch length conflicts. We used the continuous and directional random walk MCMC models, each of which was run for 10,100,000 iterations with a stepping stone sampler using 200 stones every 1,000 iterations. Burnin was set to 1,000,000 iterations. Analyses were visualized in Tracer v1.7.2 to ensure adequate effective sample size, mixing, and convergence. The continuous and directional models were compared using the log Bayes factor. We then reconstructed ancestral states using the better fitting model (continuous versus directional). To test if carbon and nitrogen breadths were correlated, we ran a phylogenetic generalized least squares analysis on the breadth data. This analysis was conducted in R using the gls function from nlme (v 3.1-157) using a Brownian motion model on the IQ-tree species phylogeny.

We examined the correlation, while taking phylogeny into account, between growth rates on individual carbon sources and carbon breadth across the subphylum. For each carbon source, we subset the data and the phylogenetic tree to include only species who grow on the specific carbon source. We then tested for correlation using a Phylogenetic Generalized Least Squares (PGLScaper 3.1-0) where lambda was estimated using a maximum likelihood framework.

We also examined the qualitative relationship between being a fast grower on a carbon substrate and carbon breadth. Within each carbon source, we divided all yeasts into fast, intermediate, or slow growers, excluding any that did not grow at all. This partitioning was done by defining the bottom (0.25) and top (0.75) quartiles of growth rates as “slow” and “fast”, respectively, regardless of carbon classification. All other yeasts were classified as “intermediate.” Across all carbon sources, we then totaled the number of fast, slow, and intermediate yeasts in each carbon classification. Species were included multiple times in the counts of fast and slow growers–once for each carbon source they grew on.

Generalists can utilize multiple different carbon substrates. We tested for co-evolution between carbon traits (growth or no growth on carbon substrates) and either carbon generalism or carbon specialism. We also reconstructed the gain and loss of each carbon trait across all measured species. We used BayesTraits v4.4.0 to model dependent and independent evolution across species. Growth and carbon class were converted into binary 0 (absent) and 1 (present) data. Trees were trimmed to include only species for which we had data. We ran both a discrete independent and a discrete dependent model using reverse jump MCMC with an exponential prior with a mean of 10. Models were run with the same parameters as the continuous breadth reconstruction listed above. To determine if the two traits were evolving independently or dependently, we compared the marginal likelihoods from the stepping stone sampler using the log Bayes factor. Better fitting dependent models with a log Bayes factor greater than 2 suggested coevolution between the two traits.

Models were then interrogated for transition rates. Zero rates were determined by examining model strings, which report the number of zero and non-zero rate parameters per step. Rate parameters with a zero rate in more than 80% of samples were set to zero. We then determined the frequency of the following scenarios: the frequency of trait gain (as opposed to loss) being higher in generalist background (compared to non-generalist), the frequency of trait gain and loss being equal in a generalist background (compared to non-generalist), and the frequency of trait loss.

We then compared these results with the rates of gain and loss for each carbon trait when assessed across all species. We tested models with either unrestricted rates for trait gain and loss with a reverse jump model, or we restricted equal rates for gain and loss. We then found the best-fitting model using the log Bayes factor. We compared the rates of gain and loss two ways. First, we identified the percentage of samples where the rate of loss was greater, less than, or equal to the rate of gain. Second, we identified models where the mean of one rate was greater or less than one standard deviation from the mean of the samples.

This analysis was repeated in BayesTraits to interrogate the co-evolution of carbon and nitrogen classifications. Classifications were converted to binary states (generalist versus non-generalist) for each carbon and nitrogen source. Each carbon binary trait was then tested against each nitrogen binary trait for a total of 4 comparisons.

### Intrinsic Factor Analyses

KOs were used to build metabolic networks for all yeasts. Each KO was associated with KEGG enzyme (EC) and reaction data using the ko00001 data and br08202 data available on the KEGG: Kyoto Encyclopedia of Genes and Genomes (KEGG) database, respectively. KOs were associated with reactions using enzymes as a connection. These associations were then used to generate binary matrices of all enzymes and reactions present within species.

We analyzed whether the overall metabolic network structure varied between generalists and specialists. To construct those networks, for each reaction, we found the input (reactants) and output (products) molecules of that reaction in KEGG. We connected reactions to each other to generate a reaction edge list by linking the input molecules of one reaction to the products of other reactions. In other words, when the product of one reaction was the input of another reaction, we created an edge between those two reactions. We used this reaction edge list to generate a KO edge list to construct networks for each species. The metabolic networks for each species were generated in R using the package igraph (v. 1.3.5). Networks were first generated from edge lists using the *graph.data.frame* function; we did not set the network to directed since many metabolic reactions are amphibolic. We were able to simplify the networks to remove any loops or duplicate edges using the *simplify* function.

To determine whether there was a difference in network structures among specialist, standard, and generalist yeasts, we calculated a set of network statistics for each species’ KEGG network. Network statistics were also calculated in R using igraph. To examine the correlation between carbon metabolism and the metabolic networks, we conducted a PGLS using carbon breadth and the various KEGG network metrics. PGLS analysis was carried out using the same methods as above.

To determine whether specific KEGG pathways were differentially represented or unique to generalists or specialists, we used KEGG enrichment analyses on our KOs. We first limited our KEGG analysis to those annotations that were variable across generalists and specialists. We next categorized KOs into those that were present or absent only in generalists or specialists. To be considered enriched in either generalists or specialists, that annotation needed to be present in all generalists and in less than 20% of specialists or vice versa. After KOs were classified, we used the R packages clusterProfile (v. 4.4.4) and enrichplot (v. 1.16.2) to perform and visualize KEGG enrichment analyses. We used the *compareCluster* function to detect KEGG enrichments. The background annotations in this analysis included all KOs present in generalists and specialists. Finally, we performed a multiple test correction using the false discovery rate (fdr) option under the pAdjustMethod. The results of this analysis were plotted using the *dotplot* function in enrichplot.

During the KEGG analysis, we identified multiple KOs that are entirely or nearly exclusive to the family Saccharomycetaceae and were subsequently removed. For example, *WHI5*, a transcriptional regulator of the cycle, was annotated using a KO (K12413) that is specific to the Saccharomycetaceae. The KO did not annotate the *WHI5* homolog that has been identified in *C. albicans* (*111*). The Saccharomycetaceae-specific KOs led to a specialist enrichment due to the large number of specialists in the group. We therefore determined that any KOs that were labeled as “Yeasts” (e.g. *Cell Cycle –Yeasts)* were not generalizable to the Saccharomycotina as a whole and were disregarded.

To identify complex patterns of metabolic presence and absence associated with carbon classification algorithm, we constructed a machine learning algorithm built by an XGBoost random forest classifier (XGBRFClassifier()) with the parameters “max_depth=12, n_estimators=100, use_label_encoder=False, eval_metric=’mlogloss’, n_jobs=8”; it was trained on 90% of the genomic data (from KEGG/Interproscan), while 10% was used for cross validation, using RepeatedStratifiedKFold from sklearn.model_selection. This analysis was used to generate accuracy measures for the algorithms, as well as their receiver operating characteristic curve (ROC), area under the ROC curve (AUC), and the cross_val_predict() function from Sci-Kit learn was used to generate the confusion matrixes. Top features were automatically generated by the XGBRFClassifier using Gini importance.

Two pathways were identified from the machine learning classification work for further analysis. The first pathway analyzed is the pentose and glucuronate interconversions pathway (KEGG map00040). The presence and absence data of the various enzymes were extracted from the KOs. The xylulokinase-encoding gene *XYL3* (step 8 Fig. 4D) was only partially annotated by KEGG. To annotate *XYL3,* we used previously published reference sequences (*57*). An hmm profile was built from these sequences and then used to search the remaining yeast protein annotations using hmmscan (hmmer v3.3.2.) These sequences were extracted, visually examined for gene fragments, aligned using mafft (v7.487) (*112*), and a gene tree was generated using IQ-TREE 2 (v2.1.2) (*105*).

The second pathway is a portion of the lysine degradation pathway (KEGG map00310) that leads to the synthesis of carnitine. For each reaction of interest, we used the KEGG annotations to assess if each species could, in principle, complete that step. The last step of the pathway (reaction 1.14.11.1) was not annotated in any yeast species. This pathway, however, was previously characterized in *C. albicans* (*62*). Using the reference sequences from *C. albicans,* we confirmed the KEGG annotations for Steps 4, 5, 6, and 7 (Fig. 4E). To annotate the final reaction (Step 7; encoded by *C. albicans* gene *BBH2*), we first identified other putative yeast Bbh2 protein sequences in the NCBI database. The remaining *BBH2* sequences were identified by using the same method as *XYL3*.

### Extrinsic Factor Analyses

To test for associations between ecology and carbon classification, we generated a formal ontology of yeast isolation environments. For strains not isolated as part of this study, isolation environments for each strain were obtained from strain databases or species descriptions. Out of the 1,154 yeasts examined, we were able to identify strain-specific isolation environments for 1,088 (94%). Strains without isolation environments were either lacking any information or had been significantly domesticated via crossing or subculturing. Isolation environments varied greatly in their information content ranging from “soil” to “gut of *Lelis* sp. (Carabidae: Coleoptera), on a basidiocarp of *Thelephora* sp.” To standardize these environments, they were added to a hierarchical ontological network of environments. We focused our analysis on the direct environment from which the yeasts had been isolated (i.e., using “gut of *Lelis* sp.*”* and not “basidiocarp of *Thelophora* sp.”). Unique isolation environments were then used to create an ontology of isolation environments using Web Protégé. Briefly, isolation environments were categorized into the following exclusive classes: animal, environmental, fungal, plant, or victuals, which we define as food or beverage products made by humans. The direct environment is the environment most directly in contact with the yeast. For example, the isolation environment “*Drosophila hibisci* on *Hibiscus heterophyllus”* was directly associated with the animal sub-class “*Drosophila hibisci”.* The ontology files are available on the Figshare repository. Machine learning classification based on environmental ontology was performed using the same methods as classification based on KEGG/InterProScan data.

To examine whether generalists or specialists were associated with specific environments, we counted the numbers of generalists and specialists in the high-level classes of environments that had the greatest number of sampled species– Victuals, Environmental, Plants, Arthropoda, and Chordata. For each environment, we determined whether specific traits were enriched for generalists and specialists using a Fisher’s Exact test. Traits included all the carbon metabolism traits, the enzymes EC:3.2.1.25 and EC:3.2.1.21, and the carnitine and pentose and glucuronate interconversion pathways. All analyses were performed in R, and the visualizations were built using *ggplot2* (v. 3.4.0).

To determine whether a trait was present more often within a specific environment than across all other environments for generalists and specialists, we used a two-sided exact binomal test. This analysis was performed using the *binom.test* function in R and visualized using *ggplot2* (v. 3.4.0).

To better understand within-environment differences between generalists and specialists, we performed a phylogenetic principal components analysis (pPCA) to account for the evolutionary non-independence of our observations while reducing dimensionality. The features used in this analysis were the absolute growth-rates on the 18 carbon sources, including glucose. A small number of growth rates were missing from our data (49 out of 3,708 growth rates), and these missing values were replaced with the mean growth rate of all yeasts on that carbon source. The pPCA was conducted in R using phytools v1.2-0 (*113*). The first four components accounted for 78% of the cumulative variance and were subsequently analyzed. We then computed the normalized Mahalanobis distance between each generalist and specialist observation across all environments. The data was then subset into each of the environments (Victuals, Environmental, Plants, Arthropoda, and Chordata), and the generalist to specialist differences were recomputed. These distances were then clustered using standard average hierarchical clustering to visualize the raw growth rates on a heatmap built using heatmap3 in R.

## Acknowledgments

We thank Linda C. Horianopoulos, Kaitlin J. Fisher, Liang Sun, David J. Krause, Kyle T. David, Christina M. Chavez, David C. Rinker, Trey K. Sato, and Hittinger Lab and Rokas Lab members for helpful discussions; the yeast community for publicly depositing taxonomic type strains; the University of Wisconsin Biotechnology Center DNA Sequencing Facility (Research Resource Identifier – RRID:SCR_017759) for providing DNA sequencing facilities and services; Wisconsin Energy Institute staff for computational support; and the Center for High-Throughput Computing at the University of Wisconsin-Madison (https://doi.org/10.21231/GNT1-HW21). This work was performed in part using resources contained within the Advanced Computing Center for research and Education at Vanderbilt University in Nashville, TN. Mention of trade names or commercial products in this publication is solely for the purpose of providing specific information and does not imply recommendation or endorsement by the US Department of Agriculture. USDA is an equal opportunity provider and employer.

## Funding

National Science Foundation Grant DEB-1442148 (CTH)

National Science Foundation Grant DEB-2110403 (CTH)

National Science Foundation Grant DEB-1442113 (AR)

National Science Foundation Grant DEB-2110404 (AR)

DOE Great Lakes Bioenergy Research Center, funded by BER Office of Science Grant DE-SC0018409 (CTH)

USDA National Institute of Food and Agriculture Hatch Project 1020204 (CTH)

H. I. Romnes Faculty Fellow, supported by the Office of the Vice Chancellor for Research and Graduate Education with funding from the Wisconsin Alumni Research Foundation (CTH)

National Institutes of Health/National Institute of Allergy and Infectious Diseases Grant R56 AI146096 (AR)

National Institutes of Health/National Institute of Allergy and Infectious Diseases Grant R01 AI153356 (AR)

Burroughs Wellcome Fund (AR)

Research supported by the National Key R&D Program of China Grant 2022YFD1401600 (XXS)

National Science Foundation for Distinguished Young Scholars of Zhejiang Province Grant LR23C140001 (XXS)

Fundamental Research Funds for the Central Universities Grant 226-2023-00021 (XXS)

National Institutes of Health Grant T32 HG002760-16 (JFW)

National Science Foundation Grant Postdoctoral Research Fellowship in Biology 1907278 (JFW)

Howard Hughes Medical Institute through the James H. Gilliam Fellowships for Advanced Study program (JLS, AR)

National Science Foundation Graduate Research Fellowship Grant DGE-1256259 (QLK)

Predoctoral Training Program in Genetics, funded by the National Institutes of Health Grant 5T32GM007133 (QLK)

Slovenian Research Agency Grant P4–0116 (NC)

Slovenian Research Agency Grant MRIC-UL ZIM, IP-0510 (NC)

Fundação para a Ciência e a Tecnologia Grant UIDP/04378/2020 (CG, PG, JPS)

Fundação para a Ciência e a Tecnologia Grant UIDB/04378/2020 (CG, PG, JPS)

Fundação para a Ciência e a Tecnologia Grant LA/P/0140/2020 (CG, PG, JPS)

Fundação para a Ciência e a Tecnologia Grant PTDC/BIA-EVL/0604/2021 (CG)

Fundação para a Ciência e a Tecnologia Grant PTDC/BIA-EVL/1100/2020 (PG)

Conselho Nacional de Desenvolvimento Científico e Tecnológico - Brazil CNPq Grant 408733/2021 (CAR)

Conselho Nacional de Desenvolvimento Científico e Tecnológico - Brazil CNPq Grant 406564/2022-1 - “INCT Yeasts: Biodiversity, preservation and biotechnological innovation”

MINCyT Grant PICT-2020-SERIE A-00226 (DL)

CONICET Grant PIP 11220200102948CO (DL)

UNComahue Grant 04/B247 (DL)

## Author contributions

DAO designed and implemented research, led phenotypic data collection, led genome sequencing, performed computational analyses and statistical analyses, managed data, and prepared the figures.

ALL designed and implemented computational analyses, managed data, and prepared figures.

MCH designed and implemented machine learning analyses.

JFW designed and implemented genome filtering and data curation pipelines.

JK and JLS. conducted data curation and filtering.

XXS. led the phylogenomic analyses with C.L. and Yuanning Li.

XZ led the annotation of genomes with Yonglin Li.

DAO, HRS, JV, CRM, QKL, EJU, and ABH phenotyped and sequenced strains.

M.S. and C.G. performed fructophilic phenotyping experiments.

DAO, KVB, MJ, MABH, QKL, CAR, NC, DL, CPK, MG, and CTH provided yeast strains.

JHD, ABH, CPK, and MG curated and organized strains and metadata.

JPS contributed resources to fructophilic phenotyping experiments.

PG supervised fructophilic phenotyping experiments.

CPK and MG led the taxonomy.

DAO and ALL co-wrote the manuscript with contributions from MCH, JFW, JK, JLS, CG, PG, XZ, XXS, MG, AR, and CTH

AR and CTH edited the manuscript.

CPK, AR, and CTH designed the research, obtained funding, and supervised the project.

All authors provided comments, input, and approved the manuscript.

## Competing interests

JLS is a scientific adviser for WittGen Biotechnologies and an adviser for ForensisGroup, Inc. AR is a scientific consultant for LifeMine Therapeutics, Inc. The other authors declare no other competing interests.

## Data and materials availability

All genome sequence assemblies and raw sequencing data have been deposited in GenBank under accessions that will be made available upon acceptance of the manuscript. All other data, including data on growth on different carbon and nitrogen sources and isolation environment data, have been deposited in Figshare (the public link will be made available upon acceptance of the manuscript). All code has been deposited in GitHub at https://github.com/DanaO523/Y1000Project. Nearly all strains came from globally recognized yeast culture collections and may be ordered from the United States Department of Agriculture (https://nrrl.ncaur.usda.gov for NRRL strains) or Westerdijk Fungal Biodiversity Institute (https://wi.knaw.nl for CBS strains) under their respective Material Transfer Agreements (MTAs) for publicly deposited strains. Strains that represent candidates for novel species that have not yet been formally described may be obtained from under the Uniform Biological MTA or other mutually acceptable MTA.

## Supplementary Figures

**fig. S1:** The phylogeny of 1,154 budding yeasts and fungal outgroups built from 2,408 orthologous groups of genes. Branches are colored according to their taxonomic assignment to an order of Saccharomycotina.

**fig. S2:** High overall congruence between concatenation and coalescent based trees of the Saccharomycotina. The highest incongruence occurred within the Serinales, which is also the largest order (420 strains). A total of 60 out of 1,153 (5%) nodes were incongruent.

**fig. S3: There is variation in carbon and nitrogen metabolism across Saccharomycotina**. A) Boxplots of growth rates of yeasts on carbon sources. B) Boxplots of growth rates of yeasts on nitrogen sources.

**fig. S4: There is variation in the breadth of carbon and nitrogen metabolism across Saccharomycotina**. A) Distribution of carbon breadth across Saccharomycotina. B) Distribution of nitrogen breadth across Saccharomycotina.

**fig. S5: Not all yeasts have their fastest growth rates on glucose**. Bar graph displaying the proportion of yeasts that are glucophilic and non-glucophilic.

**fig. S6: Four out of six KEGG network statistics were not significantly or positively correlated with carbon breadth.** Using a PGLS, we found no correlation between carbon breadth and network diameter (p-value=0.51), network density (p-value=0.53), network betweenness (p-value=0.49), and network modularity (0.65).

**fig. S7: Machine learning fails to accurately classify generalists and specialists based on environmental information.** The isolation environment ontology was used to train a random forest algorithm to classify generalists and specialists. The resulting algorithm was only slightly better than random at classifying these species.

**fig. S8: Phylogenetic PCA of growth rate data distinguishes between carbon generalists and carbon specialists**. Distances on PC1-4 were calculated based on this PCA to identify specific isolation environments in which generalists and specialists were more, or less, similar than in the overall dataset. Figure 5D includes subsets of the PCA shown here.

## Supplementary Tables

**table S1A & S1B**: Taxonomy, strain ID, source, and genomic information for all Saccharomycotina strains and the outgroups.

**table S2**: Classifications of specialist, standard, and generalist yeasts for all species with genomic and phyenotypic data and the growth rates of those species on the suite of carbon and nitrogen sources tested. Growth rates were calculated in the R package *grofit* (v. 1.1.1-1).

**table S3:** Results from fructophilic growth data performed by a second lab.

**table S4:** Classification of each budding yeast species as a fast, intermediate, or slow grower on each carbon source.

**table S5:** PGLS analysis of carbon breadth versus growth rate. Adjusted p-values were determined using the Benjamini-Hochberg method.

**table S6:** PGLS analysis of the relationship between carbon breadth and nitrogen breadth.

**table S7:** Analysis of reverse jump Bayes Traits. Relative gain and loss of the carbon trait was measured either in the generalist background or across the whole tree.

**table S8:** Network statistics for all yeast species.

**table S9:** KEGG enrichment analysis for annotations enriched in generalists and depleted in specialists.

## Notes

### Competing Interest Statement

J.L.S. is a scientific adviser for WittGen Biotechnologies and an adviser for ForensisGroup, Inc. A.R. is a scientific consultant for LifeMine Therapeutics, Inc. The other authors declare no other competing interests.

### Summary of Updates

accession numbers

