## Supplementary material for "Genomic and ecological factors shaping specialism and generalism across an entire subphylum": Figures S1-S8

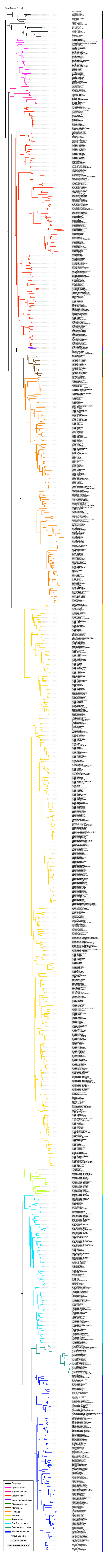

Time Scale: 0.12
Outgroup
Lipomycetales
Trigonopsidales
Dipodascales
Sporopachydermiales
Alloascoideales
Alaninales
Pichiales
Serinales
Ascoideales
Phaffomycetales
Saccharomycodales
Saccharomycetales
Public Genome
Y1000+ Genome
New Y1000+ Genome

**fig. S1**: The phylogeny of 1,154 budding yeasts and fungal outgroups built from 2,408 orthologous groups of genes. Branches are colored according to their taxonomic assignment to an order of Saccharomycotina.

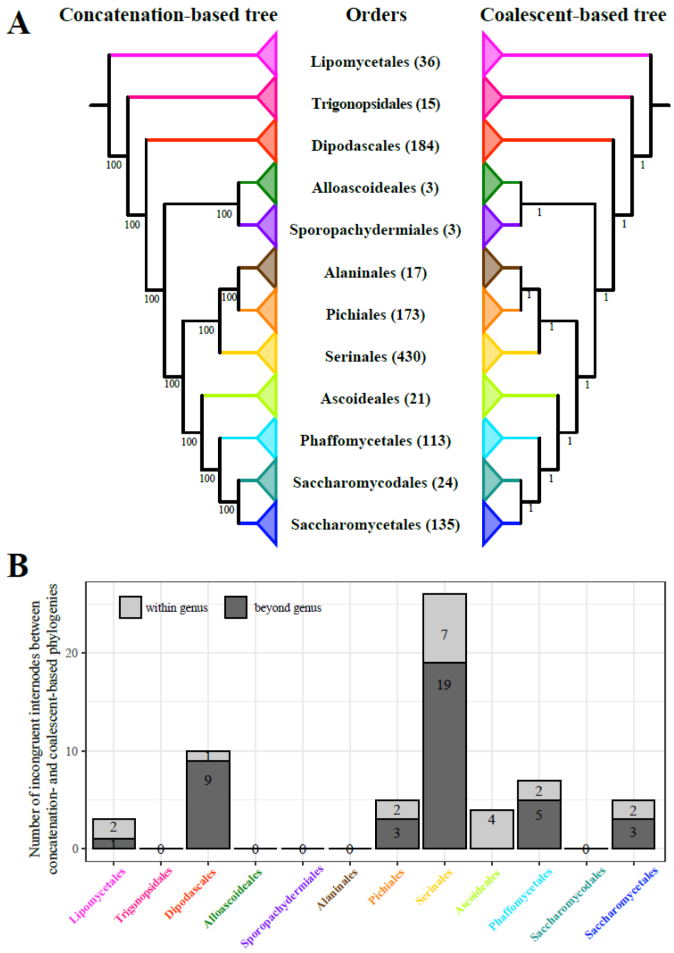


**fig. S2:** High overall congruence between concatenation and coalescent based trees of the Saccharomycotina. The highest incongruence occurred within the Serinales, which is also the largest order (420 strains). A total of 60 out of 1,153 (5%) nodes were incongruent.

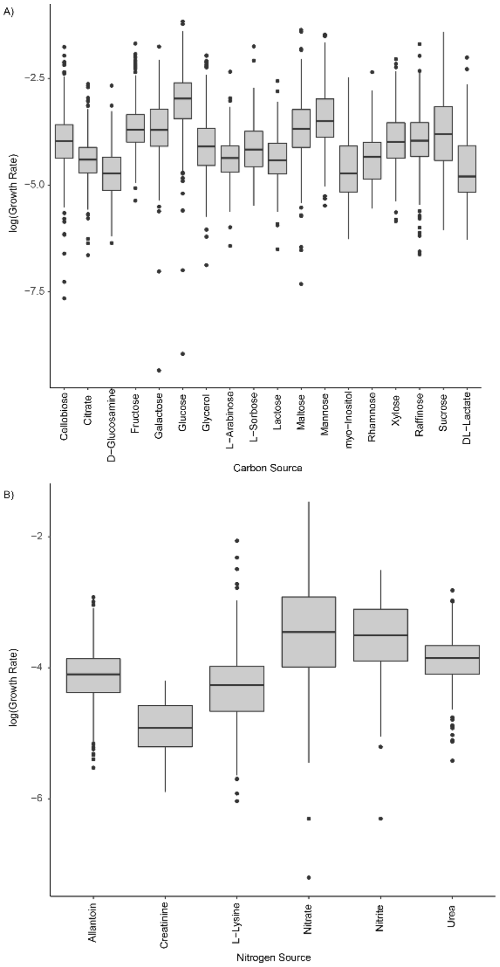


**fig. S3: There is variation in carbon and nitrogen metabolism across Saccharomycotina**. A) Boxplots of growth rates of yeasts on carbon sources. B) Boxplots of growth rates of yeasts on nitrogen sources.

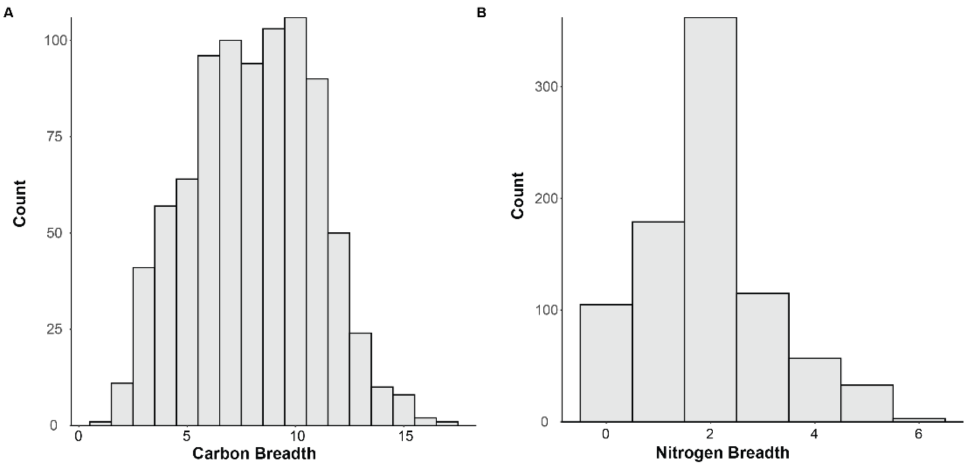


**fig. S4: There is variation in the breadth of carbon and nitrogen metabolism across Saccharomycotina**. A) Distribution of carbon breadth across Saccharomycotina. B) Distribution of nitrogen breadth across Saccharomycotina.

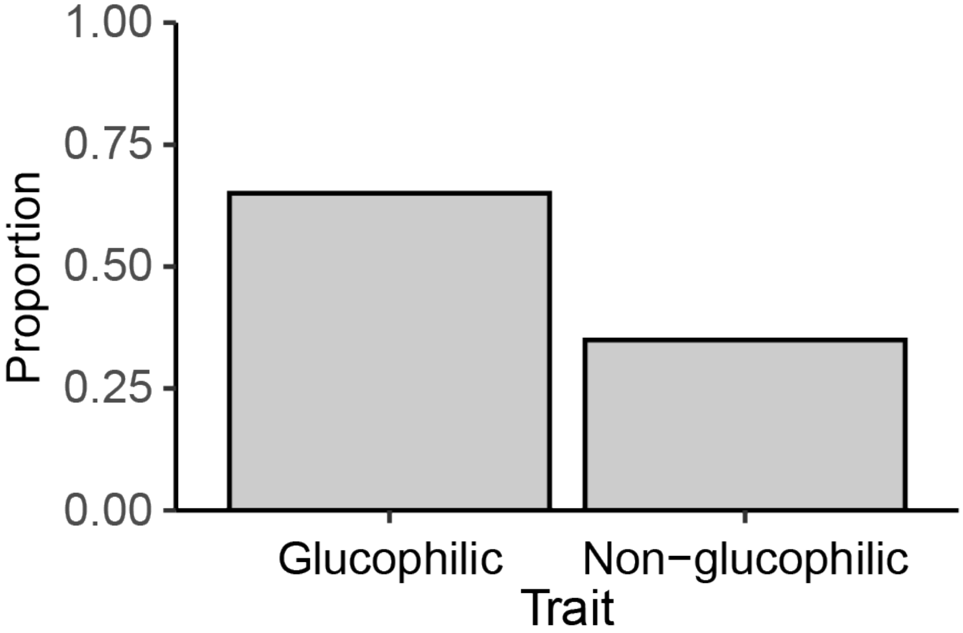


**fig. S5: Not all yeasts have their fastest growth rates on glucose**. Bar graph displaying the proportion of yeasts that are glucophilic and non-glucophilic.

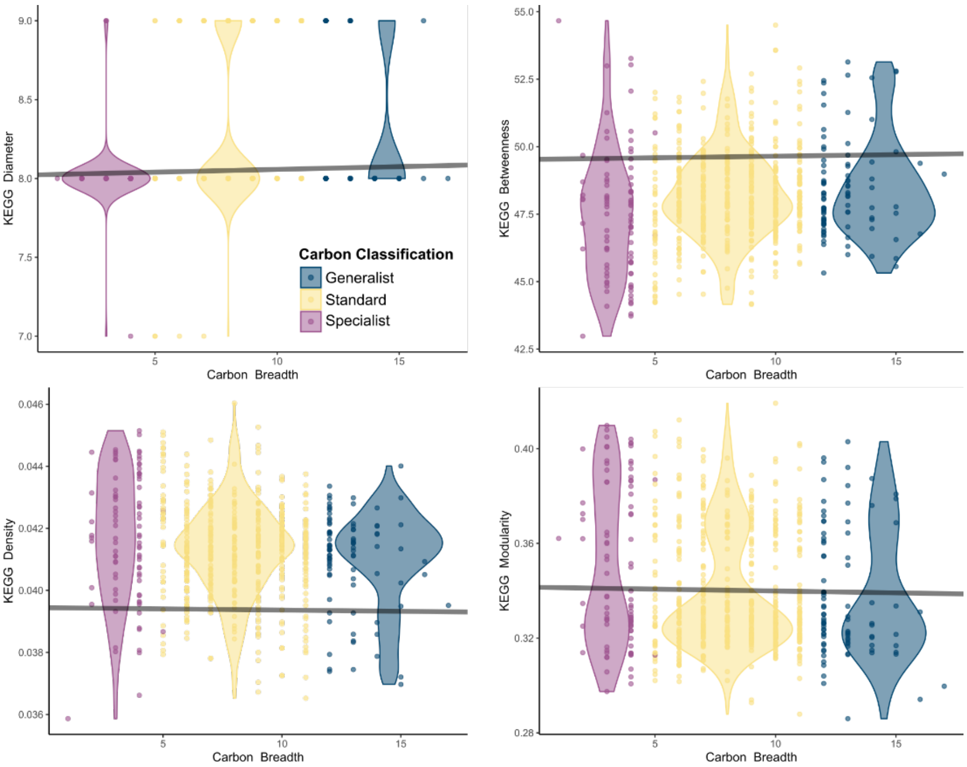


**fig. S6: Four out of six KEGG network statistics were not significantly or positively correlated with carbon breadth.** Using a PGLS, we found no correlation between carbon breadth and network diameter (p-value=0.51), network density (p-value=0.53), network betweenness (p-value=0.49), and network modularity (0.65).

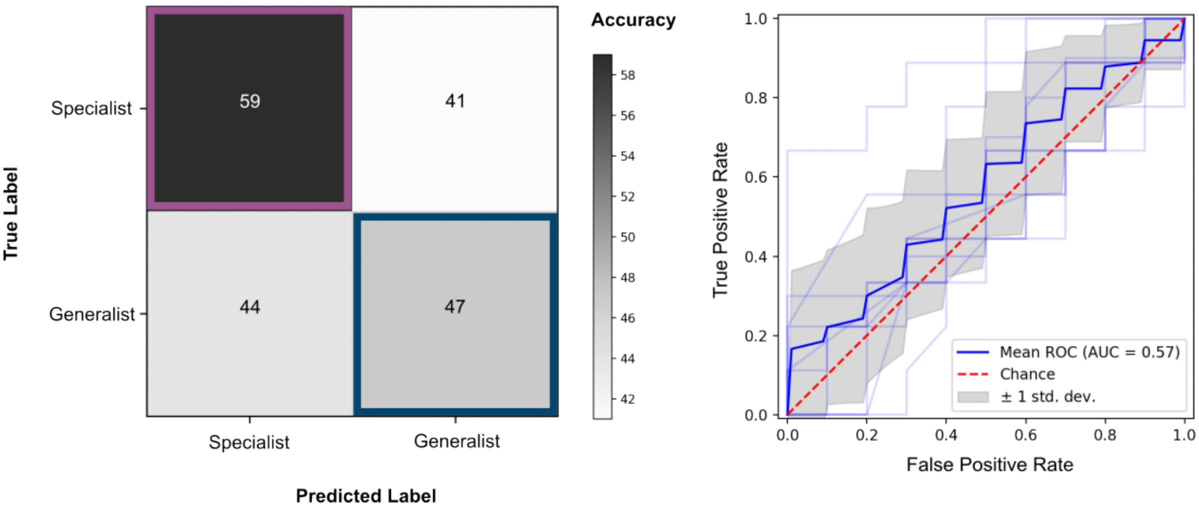


**fig. S7: Machine learning fails to accurately classify generalists and specialists based on environmental information.** The isolation environment ontology was used to train a random forest algorithm to classify generalists and specialists. The resulting algorithm was only slightly better than random at classifying these species.

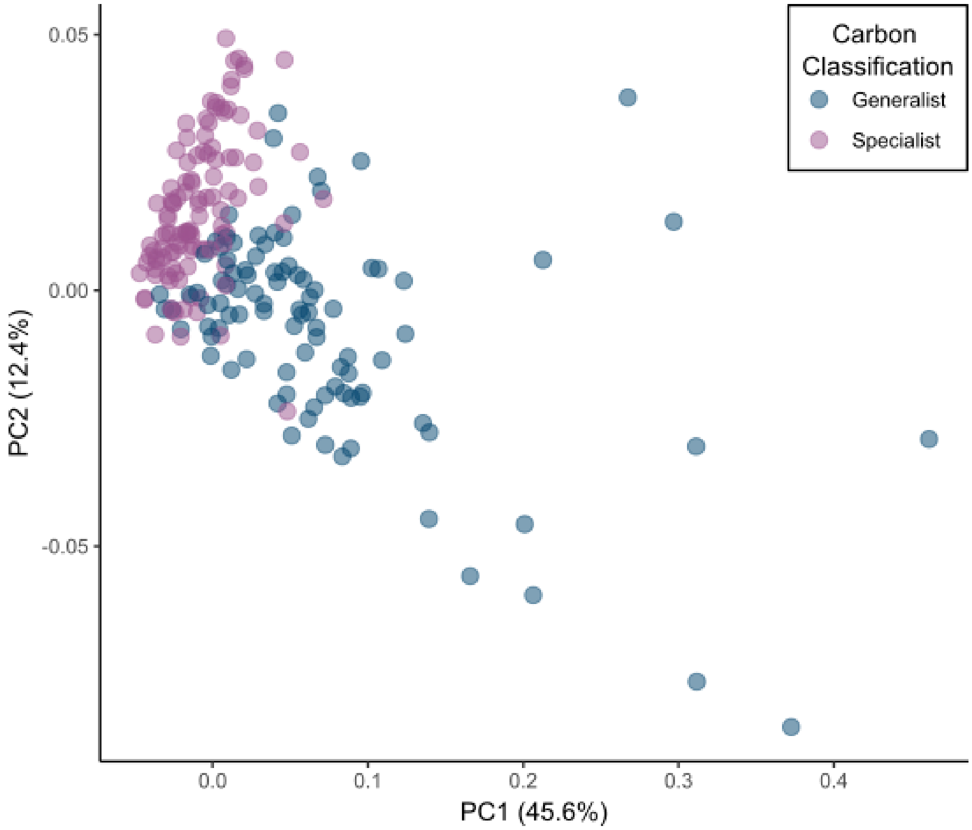


**fig. S8: Phylogenetic PCA of growth rate data distinguishes between carbon generalists and carbon specialists**. Distances on PC1-4 were calculated based on this PCA to identify specific isolation environments in which generalists and specialists were more, or less, similar than in the overall dataset. Figure 5D includes subsets of the PCA shown here.
